# Neuroprotective action of agonists and modulators of A_1_ adenosine receptors upon hyperexcitation: mechanism of the antiepileptic activity and role of neuron-glial interaction

**DOI:** 10.1101/2025.10.21.683073

**Authors:** Sergei G. Gaidin, Sergei A. Maiorov, Denis P. Laryushkin, Kristina A. Kritskaya, Alexey V. Berezhnov, Valentina N. Mal’tseva, Polina E. Rhazantseva, Irina A. Gorbunova, Daria A. Shipilovskikh, Yulia O. Sharavyeva, Sergei A. Shipilovskikh, Ivan A. Andreev, Nina K. Ratmanova, Olga A. Ivanova, Igor V. Trushkov, Danila Y. Apushkin, Alexander I. Andreev, Eugenia A. Ahremenko, Artem M. Kosenkov

## Abstract

Hyperexcitation of neuronal networks is believed to be the main reason for the excitotoxic death of neurons in different central nervous system pathologies, including epilepsy, ischemic stroke, and traumatic brain injury. G_i_-coupled receptors can be considered as promising targets for the development of new neuroprotectors. Here, we studied the anticonvulsant activity of the agonists and positive allosteric modulators (PAM) of A_1_ adenosine receptors (A_1_Rs). Our experiments demonstrate that A_1_R agonists, CCPA and N^6^-cyclohexyladenosine (N^6^-CHA), suppress hyperexcitation in three different *in vitro* models, including acute glutamate excitotoxicity, NH_4_Cl– and bicuculline-induced epileptiform activity. We have found that the inhibitory action of the agonists is mediated by the activation of not only the neuronal A_1_Rs but also the astrocytic receptors. In astrocytes, A_1_R agonists enhance GABA release, possibly via induction of calcium transients. Using inhibitory analysis, we have demonstrated that Gβγ-mediated activation of phospholipase C and subsequent Ca^2+^ mobilization from internal stores are essential for generating calcium transients in astrocytes following N^6^-CHA application. We have shown first that Ca^2+^-dependent activation of protein kinase C, which is involved in the mechanism of GABA release by astrocytes, is a pivotal step in the realization of the antiepileptic action of A_1_R agonists. Moreover, using the model of epileptiform activity induced by GABA_A_R blockade, we have shown that PAMs, PD81723 and VCP171, also suppress hyperexcitation. Furthermore, using the picrotoxin-induced epilepsy model in mice, we demonstrated that A_1_R agonists exhibit significant anticonvulsant effects and improve animal survival. The PAMs PD81723 and VCP171, when administered one hour before seizure induction, did not significantly affect seizure severity or survival rates. However, chronic administration of VCP171 produced a pronounced anticonvulsant effect and significantly increased survival.. Importantly, PAMs provided therapeutic benefits without significantly affecting overall activity levels in mice. Thus, our study demonstrates that both agonists and PAMs of A_1_R can be considered as potential therapeutic agents with antiepileptic and neuroprotective activity.

## 1. Introduction

Hyperexcitation of neuronal networks accompanies various central nervous system (CNS) pathologies, including epilepsy, ischemic stroke, and traumatic brain injuries (TBI). In all cases, a shift of the inhibition/excitation balance towards excitation is the underlying cause. In epilepsy, this can occur due to congenital reasons or as a result of acquired disorders. At the same time, in the case of ischemic stroke and traumatic brain injuries, hyperexcitation emerges from the excessive accumulation in the extracellular space of excitatory mediators (excitotoxicity), such as glutamate and aspartate, which occurs as a result of the massive death of nerve cells, followed by loss of their plasmatic membrane integrity ^1^. In turn, hyperexcitation promotes neuronal damage and an increase in lesion focus. A sustained rise in intracellular Ca^2+^ concentration (Ca^2+^ overload), leading to subsequent mitochondrial dysfunction, oxidative stress, and impairment of membrane permeability, is considered one of the primary causes of apoptotic or necrotic neuronal death upon excitotoxicity ^2^. As recent studies demonstrate, GABAergic neurons are more susceptible to excitotoxic damage due to their more pronounced excitability resulting from insufficient inhibitory innervation compared to glutamatergic neurons ^3,4^. The loss of GABAergic neurons promotes the formation of abnormal neuronal networks where the E/I balance is shifted towards excitation. These changes in the neuronal network architecture underlie the development of post-traumatic epilepsy in patients who have suffered a stroke or TBI ^3^. The absence of effective neuroprotective drugs and the high incidence of drug-resistant epilepsy (up to 30% of cases) make the search for new approaches to the treatment of these pathologies a crucial scientific task.

G-protein-coupled receptors (GPCRs), particularly those coupled to Gi protein, can be promising targets for developing new therapeutic approaches. The members of the GPCR superfamily share a similar molecular architecture, comprising seven α-helices that span the membrane, an extracellular N-terminus with N-glycosylated sites for receptor transport, and an intracellular C-terminus with phosphorylation sites for regulating desensitization. The G-protein is a heterotrimeric complex composed of a Gα-subunit bound to a dimer of Gβγ-subunits. Since Gβγ-subunits are quite conservative, GPCR classification is based on their α-subunits, which are divided into Gαs, Gαi/o, Gαq/11, Gα12/13. Receptors associated with the Gαi/o-subunit are of particular interest in the context of suppressing hyperexcitation, as their activation generally leads to inhibition of adenylyl cyclase and attenuation of neuronal excitability ^5^. G_i_-protein-coupled receptors with a significant representation in the brain include adenosine A_1_ and A_3_, cannabinoid CB_1_ and CB_2_, GABA_B_ receptors, serotonin 5-HT_1_ receptors, as well as metabotropic glutamate receptors of groups II and III. This work focuses on A_1_ adenosine receptors (A_1_Rs) since their activation, as we have previously shown ^6^, suppresses and even prevents the development of epileptiform activity *in vitro*.

Adenosine synthesis occurs with the participation of membrane and cytosolic 5’-nucleotidases CD39 and CD73, which catalyze the hydrolysis of ATP, ADP, and AMP to adenosine. Its degradation is carried out with the participation of adenosine deaminase, which catalyzes the conversion of adenosine to inosine ^7^. These enzymes maintain the concentration of adenosine in the brain at an average of 30-300 nM ^7,8^. Adenosine can activate four subtypes of its G-protein-coupled receptors: A_1_, A_2A_, A_2B_, and A_3_. The downstream intracellular effects of their activation vary widely depending on the cell type. A_1_Rs are most prevalent in the brain, with A_2B_ and A_3_ receptors (A_2B_Rs and A_3_Rs) less represented. A_2A_ receptors (A_2A_Rs) are localized predominantly in the striatum, olfactory tubercle, and nucleus accumbens ^9^. The receptors differ in their affinity for adenosine and can be ranked as follows: A_1_Rs – highest affinity (70 nM), A_2A_Rs – low affinity (150 nM), A_2B_Rs and A_3_Rs – very low affinity (5100 and 6500 nM, respectively) ^10^. A_1_Rs are widely expressed on neurons of the cortex, hippocampus, and cerebellum, as well as on astrocytes, oligodendrocytes, and microglia ^9^. In neurons, they are localized predominantly in synaptic regions, where they modulate the release of glutamate, acetylcholine, serotonin, and GABA.

Adenosine acts as an endogenous anticonvulsant suppressing neuronal excitability through the activation of both post– and presynaptic A_1_Rs. It is believed that enhanced adenosine production contributes to the termination of the seizure. The concentration of endogenous adenosine increases significantly during epileptic seizures ^8^, and according to studies performed in humans and animals, this increase occurs even before the seizure ends ^11^. A_1_R agonists also exhibit an antiepileptic effect, as demonstrated in cellular and animal models of epileptiform activity ^7,12^. In turn, genetic knockout of the A_1_R leads to lethal consequences in mice subjected to kainic acid or TBI ^13^. Despite numerous studies indicating the anticonvulsant effect of A_1_R activation, the underlying molecular mechanism remains practically unexplored. It is generally believed that the suppression of hyperexcitation is mediated by neuronal A_1_Rs, while receptors expressed on other cell types, such as astrocytes, are virtually not considered. We have previously shown that activation of α_2_-adrenergic receptors (α_2_-ARs), which are also coupled to Gi-protein, leads to an increase in the intracellular concentration of calcium ions ([Ca^2+^]_i_) in astrocytes, followed by GABA release ^14^. In these glial cells, the α_2_-ARs-mediated increase in [Ca^2+^]_i_ occurs due to the activation of the G_i_βγ-subunit and subsequent mobilization of Ca^2+^ from internal stores. Hence, this work aimed not only to study the effect of A_1_R activation in various models of hyperexcitation but also to determine whether the astrocytes are involved in the realization of the antiepileptic effect of A_1_R agonists.

To date, considerable evidence has accumulated indicating that A_1_Rs can serve as targets for the development of anticonvulsant drugs; however, the use of direct GPCR agonists is associated with various side effects, primarily in the cardiovascular system, particularly in the case of A_1_Rs. Additionally, the effectiveness of direct GPCR agonists decreases over time due to receptor desensitization ^15^. For these reasons, no drug based on a direct agonist of Gi-protein-coupled receptors has passed all stages of clinical trials. Positive allosteric modulators (PAMs) are believed to minimize side effects while maintaining positive action. Moreover, as mentioned above, in epilepsy and other pathologies, the concentration of adenosine in the CNS increases, which also creates prerequisites for minimizing side effects. This suggests that the use of A_1_R PAM can be considered an alternative approach for the development of drugs based on A_1_R ligands. In this regard, we have tested the ability of commercially available and newly synthesized A_1_R PAMs to suppress epileptiform activity.

## 2. Materials and methods

### 2.1. Preparation of mixed hippocampal neuron-glial cultures

Neuron-glial cultures were prepared from the hippocampus of newborn Wistar rats (P0-2). The hippocampus was extracted, minced using scissors, and placed in a cold Versen’s solution free of Ca^2+^ and Mg^2+^. After this, the fragments of minced tissue were incubated with a 1% trypsin solution (trypsin was diluted in Versen’s solution) for 10 minutes at 37 °C with constant mixing in a thermoshaker at 500 rpm. Then, the tissue fragments were washed twice with cold Neurobasal medium to inactivate the trypsin, and they were carefully triturated in a fresh portion of room-temperature Neurobasal medium using a pipette. The remaining large tissue fragments were removed, and the suspension was centrifuged for 3 minutes at 2000 rpm (200g). Then, the supernatant was removed, and the cells were resuspended in a growth medium consisting of Neurobasal medium supplemented with B-27 (2%) and freshly prepared glutamine (0.5 mM). Penicillin and streptomycin were also added to the growth medium. The suspension was distributed into glass cylinders (100 μL per cylinder; the inner diameter of the cylinders was 6 mm) and placed on coverslips that had been pre-treated with polyethyleneimine. Petri dishes with coverslips were placed in a CO_2_ incubator for 1 hour for cell sedimentation and adhesion. After this, the cylinders were removed, and 2 mL of growth medium was added to the dishes with coverslips. The cultures were grown in a CO_2_ incubator at 37°C and 95% humidity for two weeks, and then used in subsequent experiments. In all experiments, cultures aged 12–14 days *in vitro* were used.

### 2.2. Fluorescence microscopy

#### 2.2.1 Calcium imaging

The dynamics of [Ca^2+^]_I_ in cells of neuron-glial cultures was assessed using the fluorescent Ca^2+^-sensitive probe Fura-2 AM. The cultures were loaded with the probe dissolved to a concentration of 3 μM in Hank’s balanced salt solution (HBSS) of the following composition: 136 mM NaCl, 3 mM KCl, 0.8 mM MgSO_4_, 1.25 mM KH_2_PO_4_, 0.35 mM Na_2_HPO_4_, 1.4 mM CaCl_2_, 10 mM glucose, and 10 mM HEPES; pH 7.35. Loading was carried out for 40 minutes at 28 °C, and then the cultures were washed three times with HBSS solution and used in experiments. Fluorescent recordings were carried out using a Leica DMI6000B fluorescence microscope equipped with a Hamamatsu C9100 monochrome CCD camera, a Leica EL6000 illuminator, an HC PL APO 20×/0.7 IMM objective, an external set of excitation filters Fura-2 (BP340/30 and BP387/15 filters), and an internal FURA-2 filter cube (dichroic mirror 72100bs, emission filter HQ 540/50m). All experiments were conducted at 28 °C in HBSS solution.

Responses of individual cells were obtained as a result of processing and analysis of image series and expressed as the ratio (340/387) of signal intensities of Fura-2 fluorescence excited in the 340 nm channel (excitation peak of the Ca^2+^-bound form of the dye) to the fluorescence intensity of the probe excited in the 387 nm channel (excitation peak of the Ca^2+^-free form). ImageJ software (NIH, Bethesda, MD, USA) was used to process the image series.

#### 2.2.2 Immunocytochemistry

Immunocytochemical staining was used to identify astrocytes and microglia in the cultures. First, the neuron-glial cultures were washed three times with phosphate-buffered saline (PBS, pH = 7.35), then fixed with freshly prepared 4% paraformaldehyde solution (in PBS) for 30 minutes at room temperature. Then, the cultures were washed three times with an ice-cold PBS solution. To increase the permeability of cell membranes for antibodies, the detergent Triton X-100 was used. The step of blocking the non-specific binding of secondary antibodies was combined with permeabilization, so the blocking solution consisted of 10% goat serum and 0.1% Triton X-100 dissolved in PBS. The cells were incubated in the blocking solution at room temperature for 30 minutes. After this, the cells were washed once with a solution containing 1% goat serum and 0.1% Triton X-100. This solution was used to dilute the primary antibodies. Glial fibrillary acidic protein (GFAP) and ionized calcium-binding adaptor molecule 1 (Iba1) were used as markers for astrocytes and microglia, respectively. The cells were incubated with primary antibodies for 12 hours at 4 °C, then washed three times with PBS solution and incubated with secondary goat antibodies to rabbit and mouse immunoglobulins. Secondary antibodies were dissolved in PBS. The cells were incubated with secondary antibodies for 90 minutes at room temperature in the dark, and then washed three times with PBS solution. Hoechst 33342 (1 μg/mL, 5 minutes) was used to visualize the nuclei. A Leica DMI6000B fluorescence microscope (Leica Microsystems, Wetzlar, Germany) was used to visualize the distribution of the antibodies.

#### 2.2.3 Vital identification of neurons and astrocytes

Identification of neurons and astrocytes was based on calcium imaging. In one group of cells, the addition of the GABA_A_R antagonist, bicuculline, and potassium chloride (KCl) causes synchronous [Ca^2+^]_i_ oscillations (calcium oscillations) and an increase in [Ca^2+^]_i_ (calcium response), respectively (Fig. 1A′ – red curves, arrows with letters b and c). In the case of KCl, calcium responses occur due to depolarization caused by an increase in the extracellular potassium ion concentration. Calcium responses are observed in other cells only when ATP is added (Fig. 1A – green curves, arrow with letter d). We have previously shown by combining vital calcium imaging and immunocytochemical staining for NeuN and GFAP (markers of neurons and astrocytes, respectively) ^14^ that cells responding to the application of potassium chloride with a [Ca^2+^]_i_ increase contain NeuN, while cells responding with an increase in [Ca^2+^]_i_ to the application of ATP contain GFAP. GABA_A_Rs blockade leads to disinhibition of the neuronal network, which causes the generation of epileptiform activity. Thus, if calcium oscillations occur in a cell in response to the application of bicuculline, such a cell is identified as a neuron. In turn, all cells that did not exhibit such activity were identified as astrocytes, as microglial Iba1^+^ cells are practically absent in the cultures, and their number can be neglected, as shown in Fig. 1B.

**Figure 1.**
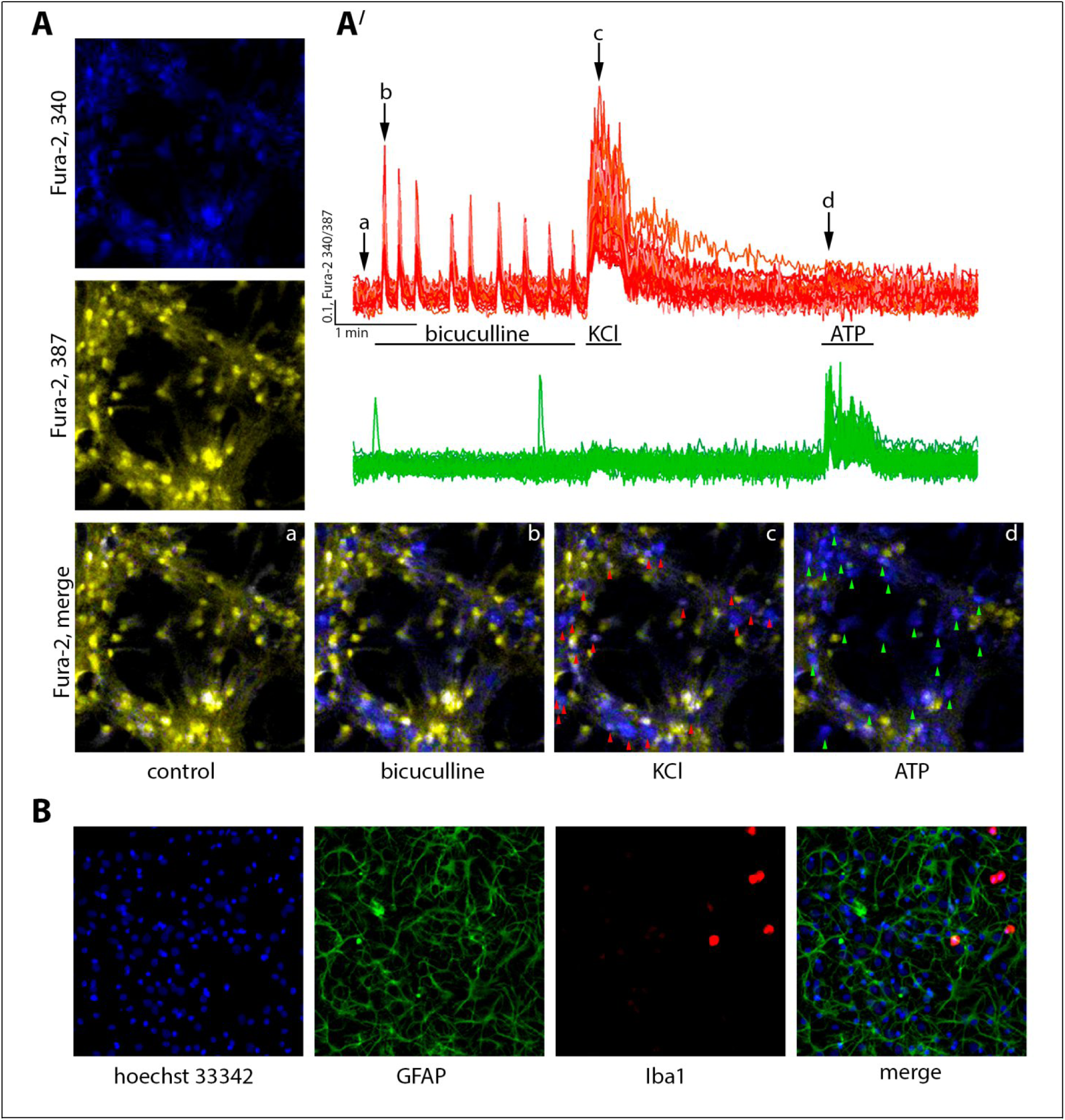
Identification of neurons and astrocytes in the hippocampal neuron-glial culture. (***A***) The images of cell cultures stained with fluorescent Ca^2+^-sensitive probe Fura-2. ***a, b, c, d*** – the images demonstrating overlay of two channels (channel with excitation 340 nm ‒ the peak of the excitation spectrum of Ca^2+^-bound form of Fura-2; channel with excitation 387 nm ‒ the peak of the excitation spectrum of Ca^2+^-free form of Fura-2) in different moments of the experiment shown in panel A′. (***A***′) Changes of intracellular Ca^2+^ concentration in neurons (red curves) and astrocytes (green curves) during the application of GABA_A_R antagonist, bicuculline (10 μM), KCl (35 mM), and ATP (10 μM). (***В***) Immunostaining of cells in the cultures with antibodies against the glial fibrillary acidic protein (GFAP) (a marker of astrocytes) and ionized calcium-binding adaptor molecule 1 (Iba1) (a marker of microglia)–Hoechst 33342 (5 μg/mL) – fluorescent probe for staining of nuclei.

### 2.3. Recording of electrophysiological activity

To record currents in neurons, the patch-clamp method was used in the “whole-cell” configuration in “voltage-clamp” mode using an Axopatch 200B amplifier (Axon Instruments, USA). Analog-to-digital conversion of data was carried out using the Axon DigiData 1440A system. Data was analyzed using the pCLAMP 10 software package (Axon Instruments, USA). All electrophysiological experiments were conducted at a temperature of 28 °C. The intracellular solution consisted of (in mM): 5 KCl, 130 K-gluconate, 1 MgCl_2_×6H_2_O, 0.25 EGTA, 4 HEPES, 2 Na_2_-ATP, 0.3 MgATP, 0.3 Na-GTP, 10 Na_2_-phospho-creatine (305-310 mOsm, pH 7.2). The composition of the extracellular solution was similar to that of the HBSS medium from section 2.2.

To detect GABA release by astrocytes, the membrane potential of the neuron was held at –30 mV. At this potential, there is an inversion of the Clgradient in neurons, resulting in chloride currents becoming outward. To exclude the contribution of neurons to GABA release, a blocker of voltage-gated sodium channels, tetrodotoxin (TTX), was used. The use of tetrodotoxin allows suppression of neurotransmission by blocking the propagation of action potentials. The Mini Analysis Program (Synaptosoft Inc., USA) was used to detect and analyze slow currents evoked by gliotransmitters.

### 2.4. Animals and housing conditions in the home cage

Male CD-1 mice weighing 28–32 g were used in this study. From the start of the experiments, the animals were housed under standard, controlled conditions (a 12-hour light/12-hour dark cycle, 20–22 °C, and 40–60% humidity) with food and water available *ad libitum*. All animal procedures were approved by the Bioethics Committee of the Institute of Cell Biophysics (ICB) and carried out according to Act708n (23 August 2010) of the Russian Federation National Ministry of Public Health, which states the rules of laboratory practice for the care and use of laboratory animals, and the Council Directive 2010/63 EU of the European Parliament on the protection of animals used for scientific purposes. ICB RAS Animal Facility provided the animals for experiments in accordance with the applications approved by by the Bioethics Commission of ICB RAS (Approval ID: 2/022025, date 2025-02-21). Throughout all experiments, animals had free access to food and water.

### 2.5. Protocol for Induction and Treatment of Epileptic Seizures

To induce seizure activity, the GABA_A_R antagonist picrotoxin was administered intraperitoneally at a dose of 5 mg/kg. Picrotoxin was dissolved in a mixture consisting of 20% (v/v) DMSO and 80% saline (0.9% NaCl, w/w). Initially, the picrotoxin powder was dissolved in anhydrous DMSO, followed by the addition of the required volume of saline. The control group received the DMSO/saline solution (20%/80% v/v) without the active compound. The primary criterion for seizure activity was the development of tonic-clonic seizures corresponding to stage 5 on the Racine scale ^16^, or stages 5–7 on the modified scale ^17^, characterized by tonic-clonic convulsions progressing to respiratory arrest and death.

Experiments were conducted in standard T3-PC cages (ZoonLab, Germany), each equipped with a video recording system (MultyNeuro-SLA, OpenScience, Moscow, Russia) and piezoacoustic sensors placed beneath the cage floor. Signals were recorded at a frequency of 992 Hz. The kinetic energy transmitted to the cage floor and walls during animal movement was converted into electrical impulses for digitization and subsequent actometric analysis. Data are presented as the per-minute envelope of the 99^th^ percentile. The recording setup included 16 individual cages, each housing one animal per experiment. Before the start of each experiment, mice were given one hour to acclimate to the new environment.

Subsequently, mice received intraperitoneal injections of the A_1_R agonists N^6^-cyclopentyladenosine (N^6^-CHA) or 2-chloro-N^6^-cyclopentyladenosine (CCPA), dissolved in the same solvent mixture as picrotoxin. Control animals received an equivalent volume of vehicle. One hour after drug administration, all mice were injected with picrotoxin. Spontaneous locomotor activity and video monitoring were conducted continuously for 24 hours following the induction. Experiments with positive allosteric modulators (PAMs) of A_1_Rs followed the same general protocol.

For PAMs (PD81723 and VCP171), a mixture of DMSO and flaxseed oil was used as the vehicle. The PAMs were first dissolved in DMSO, then mixed with flaxseed oil, and vortexed thoroughly for 3 minutes. The final concentration of DMSO in the DMSO/oil mixture was 2.5%. These compounds were also administered intraperitoneally one hour before picrotoxin injection.

In a separate series of experiments involving chronic administration of VCP171, animals were individually housed for five consecutive days. The experimental group received daily subcutaneous injections of VCP171 at a dose of 10 mg/kg, dissolved in DMSO/flaxseed oil (5 injections total). The control group received equivalent volumes of DMSO/flaxseed oil mixture. The final injection was administered one hour before the picrotoxin challenge. Animals were placed into the recording system one hour before this final injection, and the remainder of the experiment followed the same protocol as the acute treatment series.

### 2.6 Statistical analysis

#### 2.6.1 Column and group comparisons

For statistical analysis of data, OriginLab Pro 2016 software (OriginLab, Northampton, Massachusetts, USA), MS Excel (Microsoft Corporation, Redmond, Washington, USA), and Prism GraphPad 8 (GraphPad Software, San Diego, California, USA) were used. Given that the sample size was n < 15, the Shapiro-Wilk normality test was used to determine whether the data conformed to the normal distribution. The Wilcoxon test was used to test the hypothesis of homogeneity of two dependent samples, where the data do not conform to the normal distribution law. In cases where samples do not conform to a normal distribution, the Kruskal-Wallis test was used for group comparison. Group comparison of samples conforming to a normal distribution was analyzed using a two-way analysis of variance (one-tailed). The significance of differences was accepted at a significance level of p < 0.05. This study was not pre-registered. No blinding or randomization was performed in this study. There were no exclusions, and no exclusion criteria were predetermined. No test for outliers was conducted. The following parameters for box plots in the figures were defined: dimensions – 75^th^ (top) and 25^th^ (bottom) percentiles; line – median; whiskers – minimal and maximal values. Sample size calculations were performed using OriginLab software, as described previously ^4^.

#### 2.6.2 Statistical Analysis of Survival

All statistical analyses were performed in R (version 4.3.1) using RStudio (version 2023.09.1+494). Survival time (in minutes), event status (0 = censored, 1 = death), and treatment group were used as input variables. Kaplan–Meier survival curves were constructed using the *survfit*() function from the survival package. Group differences were assessed by the log-rank test, and the resulting p-values were reported. Survival curves with confidence intervals and p-values were visualized using the *ggsurvplot*() function from the *survminer* package. To estimate hazard ratios (HR) for each treatment group compared to control, we applied Firth’s penalized likelihood Cox regression using the *coxphf*() function from the *coxphf* package. This method is particularly robust in cases of small sample sizes or complete separation. Estimated HRs and their 95% confidence intervals were summarized and visualized using forest plots, generated with the *forest*() function from the meta package. A random-effects model with Hartung–Knapp adjustment was applied to account for between-group heterogeneity (method.random.ci = “HK”). All data preprocessing, group comparisons, and visualizations were performed using the following R packages: *survival*, *survminer*, *coxphf*, *meta*, and *dplyr*.

### 2.7 Chemical synthesis and characterization of the obtained substances

In this study, two groups of compounds (abbreviated as SGA and TT) with aminothiophene scaffolds similar to those in the structures of commercially available positive allosteric A_1_R modulators, PD81723 and VCP171, were synthesized. The detailed schemes of synthesis and characterization procedures for the most effective substances are provided in the Supplementary section (please see the Supporting Information file).

## 3. Results

### 3.1. The effect of A_1_R activation on neuronal activity in different models of hyperexcitation

As the first model of hyperexcitation, we used a model of epileptiform activity induced by blocking GABA_A_R (see section 2.2.3). The addition of the receptor antagonist, bicuculline (10 μM), causes synchronous calcium oscillations in neurons (Fig. 2A). The application of A_1_R agonists CCPA and N^6^-CHA causes a dose-dependent decrease in the frequency of calcium oscillations in all neurons. The maximum effect, manifesting as a complete suppression of calcium oscillations in neurons, was observed at a concentration of 100 nM for both agonists (Fig. 2A′). Using the selective A_1_R antagonist PSB36, we showed that CCPA and N^6^-CHA exert their inhibitory effect through A_1_Rs (Fig. 2B, B′).

**Figure 2.**
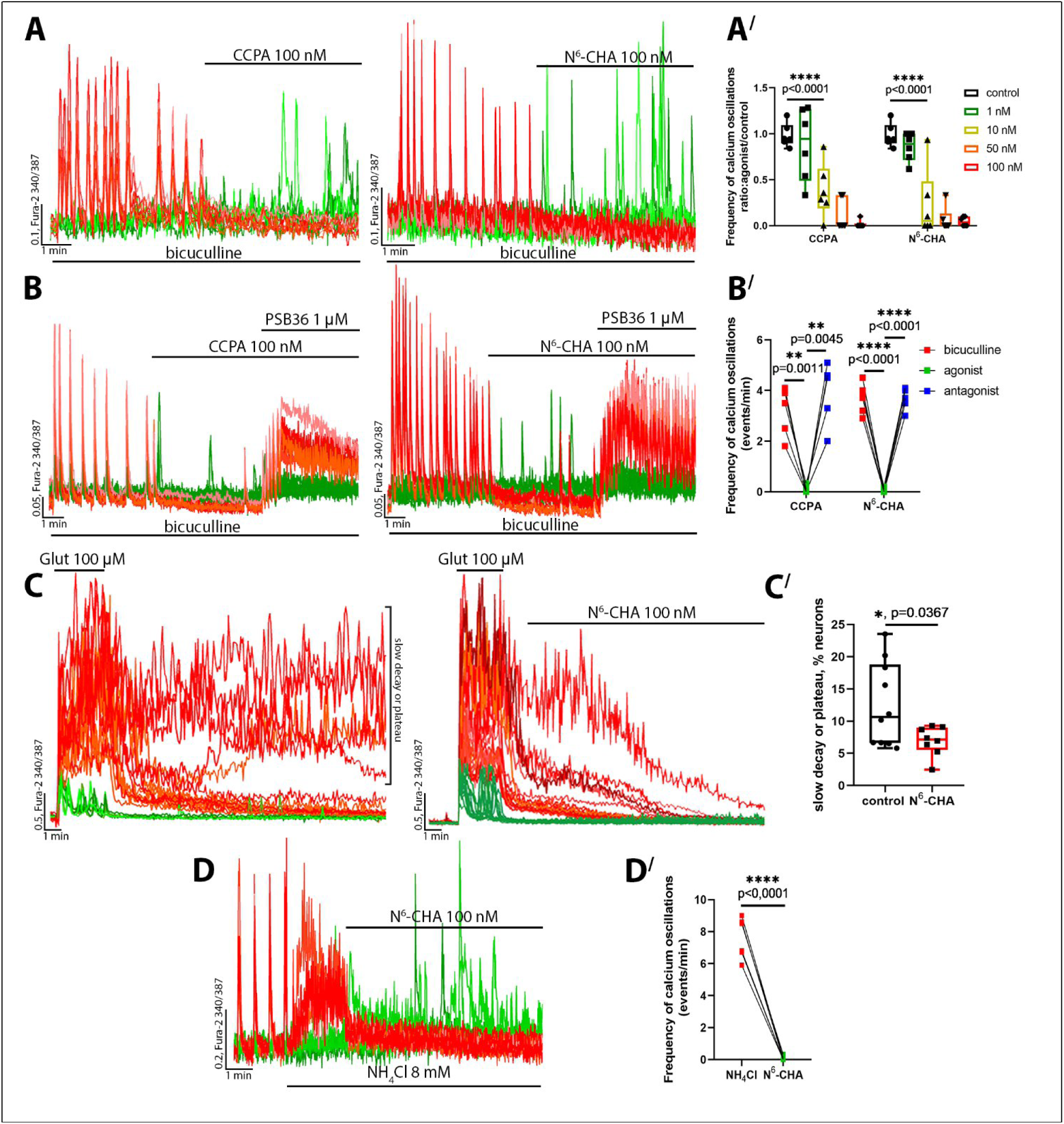
The effects of A_1_Rs activation on the neuronal network activity upon hyperexcitation. (*A*) The influence of A_1_R agonists, CCPA (100 nM) and N^6^-CHA (100 nM), on the epileptiform activity in neurons induced with the GABA_A_R antagonist, bicuculline (10 μM). (***A***′) The diagrams show the effect of the A_1_R agonists on the frequency of calcium oscillations in neurons. Ordinary two-way ANOVA followed by Sidak’s multiple comparison test; (***B***) Application of A_1_R antagonist, PSB36 (1 μM), on the background of the A_1_R agonist upon bicuculline-induced epileptiform activity. (***B***′) The diagram illustrates the effect of A_1_R agonists on the frequency of bicuculline-induced calcium oscillations in the presence of PSB36. Two-way ANOVA (across row matching) followed by Sidak’s multiple comparison test; (***C***) The effect of N^6^-CHA on the [Ca^2+^]_i_ restoration dynamics in neurons after acute glutamate excitotoxicity (application of 100 μM glutamate during 3 minutes). (***С***′) The percentage of neurons with a slow [Ca^2+^]_i_ decrease or irreversible dysregulation of the calcium homeostasis ([Ca^2+^]_i_ remains on the plateau) 20 minutes after glutamate washout. The dynamics of [Ca^2+^]_i_ restoration were considered slow if the [Ca^2+^]_i_ level was ≥ 20% of the maximal value observed during glutamate exposure. Unpaired t-test; (***D***) The effect of N^6^-CHA (100 nM) on the neuronal activity upon NH_4_Cl-induced hyperexcitation. (***D***′) The diagram shows the frequency of NH_4_Cl-induced calcium oscillations in neurons before and after N^6^-CHA application. Paired t-test. In panels A–D, red curves correspond to the responses of representative neurons, while green curves correspond to the responses of astrocytes. The number of random neurons and astrocytes analyzed in each experiment was 100 for both cell types. The dots in diagrams correspond to the mean value of the parameter in an individual experiment.

In turn, the application of A_1_R agonist increased the activity of astrocytes, as evidenced by the generation of calcium transients (Fig. 1A, green curves). Since the effects of CCPA and N^6^-CHA are similar and are realized through A_1_Rs, as confirmed by inhibitory analysis, we used only N^6^-CHA at a concentration of 100 nM in further experiments.

Glutamate excitotoxicity was used as a second model of hyperexcitation. As shown in Figure 2C, a high dose of glutamate evokes a rapid rise in [Ca^2+^]_i_ in neurons, which persists for as long as the application lasts (3 minutes). After glutamate washout with HBSS, in most neurons, the [Ca^2+^]_i_ level recovers fairly quickly; however, in approximately 11% (median value 10.64%) of neurons, the [Ca^2+^]_i_ level does not return to the baseline value even 20 minutes after glutamate washout. The application of N^6^-CHA 2 minutes after glutamate removal promotes the restoration of calcium homeostasis, as evidenced by the decrease in the percentage of neurons with high [Ca^2+^]_i_ levels to 7% (median value: 7.16) (Fig. 2C′). A slight increase in [Ca^2+^]_i_ level during glutamate application is also observed in astrocytes, but intracellular Ca^2+^ concentration rapidly decreases in these cells after glutamate removal.

In the NH_4_Cl-induced (8 mM) hyperexcitation model, activation of A_1_Rs also has a positive effect. As shown in Figure 2D, the application of ammonium chloride causes a [Ca^2+^]_i_ increase in neurons accompanied by calcium oscillations. N^6^-CHA rapidly reduces the [Ca^2+^]_i_ level in all neurons and suppresses calcium oscillations. At the same time, as in the case of experiments with bicuculline, the frequency of calcium transient generation in astrocytes increases after the addition of the agonist.

Thus, based on the experiments performed, it can be concluded that activation of A_1_Rs upon induced hyperexcitation of neuronal networks contributes to the suppression of epileptiform activity and normalization of calcium homeostasis. Nevertheless, it should be noted that the suppression of excessive induced activity in neurons is accompanied, on the contrary, by an increase in astrocyte activity.

### 3.2. The mechanism of antiepileptic activity of A_1_R agonists

To study the mechanism of the inhibitory action of A_1_R agonists, we used a model of epileptiform activity induced by blocking GABA_A_R. We tested different inhibitors of intracellular signaling pathways and a Ca^2+^-activated K^+^ (SK) channel blocker. Pharmacological analysis showed that inhibitors of βγ-mediated signaling (gallein, 50 μM), phospholipase C (U73122, 5 μM), and protein kinase C (Gö 6976, 5 μM) significantly attenuated the inhibitory action of N^6^-CHA against the bicuculline-induced calcium oscillations in neurons (Fig. 3B, C, E, G). Although the frequency of the calcium oscillations decreased after the N^6^-CHA application in the presence of each of these inhibitors, we did not observe the complete suppression of the oscillations as in control experiments with 100 nM of N^6^-CHA (Fig. 3A). In turn, the calmodulin antagonist (calmidazolium, 5 μM) and the blocker of SK channels (apamin, 100 nM) did not affect the inhibitory action of N^6^-CHA (Fig. 3Е, F). Moreover, gallein and U73122 abolished the effect of N^6^-CHA on astrocyte activity (Fig. 3H). In control experiments, N^6^-CHA application increased the number of astrocytes generating calcium transients and the number of transients occurring in a single astrocyte; however, this trend was not observed in the presence of gallein and U73122.

**Figure 3.**
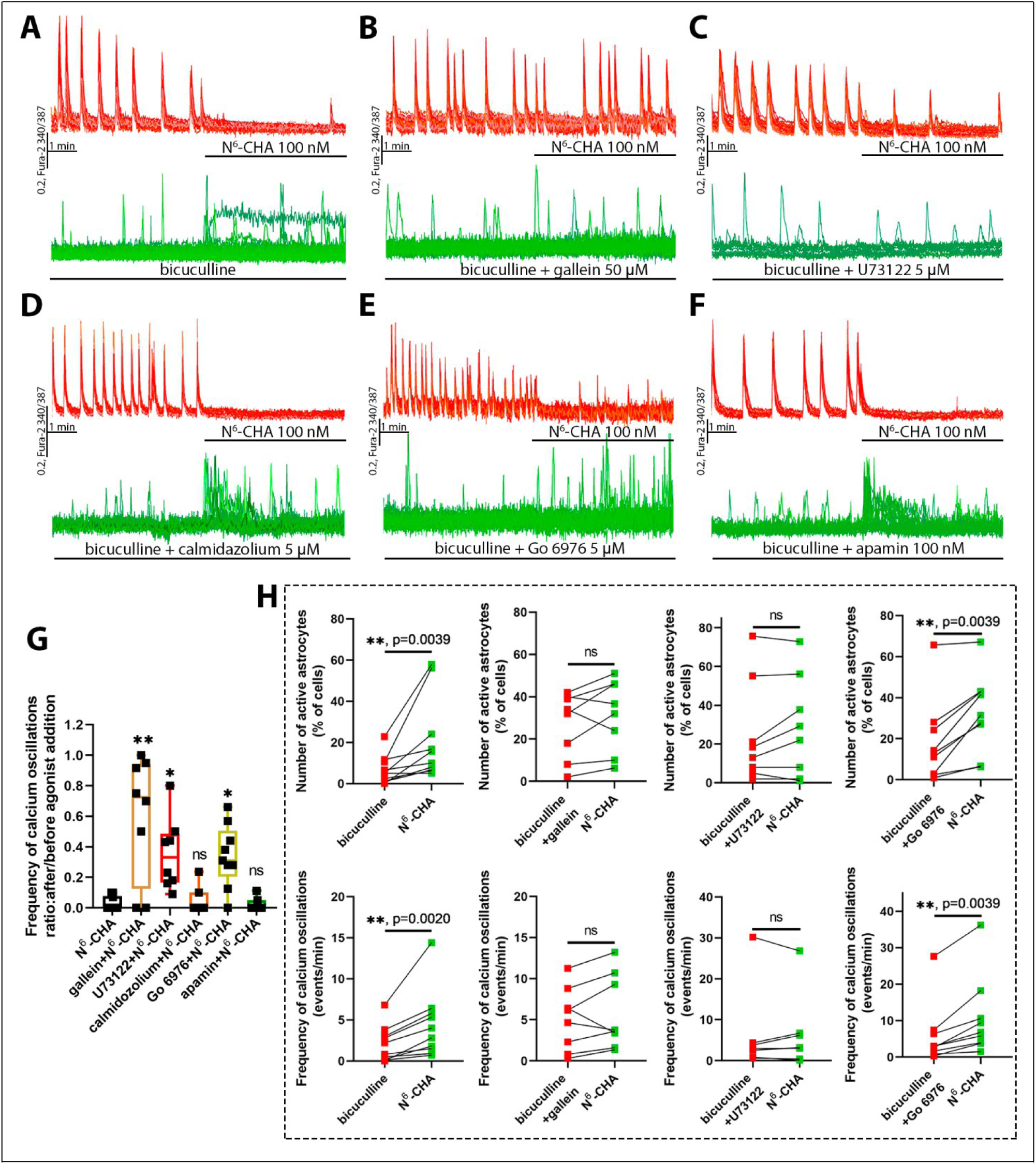
The mechanism of antiepileptic action of A_1_R agonist. (***A***) The effect of A_1_R agonist, N^6^-CHA, on epileptiform activity in neurons induced by GABA_A_R antagonist, bicuculline (10 μM) in control and in the presence of inhibitors of βγ-signaling (***B***) (gallein 50 μM, preincubation 30 minutes), PLC (***C***) (U73122 5 μM, preincubation 30 minutes), calmodulin (***D***) (calmidazolium 5 μM, preincubation 30 minutes), (***E***) PKC (Go 6976 5 μM, preincubation 30 minutes) and a blocker of SK channels (***F***) (apamin 100 nM). Red curves on the top of each panel ‒ the changes of [Ca^2+^]_i_ in representative neurons; green curves on the bottom of panels – [Ca^2+^]_i_ changes in representative astrocytes in the same experiment. We analyzed 100 randomly chosen neurons and 100 astrocytes in each experiment. (***G***) The diagram illustrates the effect of inhibitors/blockers on the frequency of calcium oscillations in neurons following the application of N^6^-CHA. The Kruskal‒Wallis test followed by Dunn’s multiple comparisons test. The significance of differences compared to control (N^6^-CHA only): gallein + N^6^-CHA ‒ ** p=0.0033; U73122 + N^6^-CHA ‒ *p=0.0122; Gö 6976 + N^6^-CHA ‒ *p=0.0152. (***H***) The diagrams show the effects of gallein, U73122, and Gö 6976 on the percentage of astrocytes responding with generation of calcium transients to N^6^-CHA application and the frequency of these transients. Wilcoxon matched-pairs signed-rank test. The dots in the diagrams correspond to the mean value of the parameter in a single experiment.

These results suggest that the Gβγ–PLC signaling cascade in astrocytes contributes to the antiepileptic action of A_1_R agonists. In turn, activation of A_1_Rs in the presence of the PKC inhibitor leads to an increase in the frequency of calcium transients in astrocytes, but does not suppress calcium oscillations in neurons (Fig. 3E, H). Therefore, as with Gβγ and PLC, PKC may be involved in the realization of the antiepileptic action of A_1_R agonists, but its recruitment occurs after [Ca^2+^]_i_ increases.

Thus, considering the obtained data and our previous studies ^14,18^, it can be assumed that the activation of astrocytic A_1_Rs induces the activation of the Gβγ–PLC signaling pathway, leading to the generation of calcium transients in astrocytes. In this case, the endoplasmic reticulum is likely a source of Ca^2+^ inflow since PLC promotes IP_3_ formation, which interacts with IP_3_ receptors, inducing Ca^2+^ mobilization from internal stores. After that, Ca^2+^-dependent activation of PKC, followed by the release of gliotransmitters, including GABA, occurs.

We used the patch-clamp technique in whole-cell mode to determine whether activation of A_1_Rs enhances GABA release by astrocytes. The release of gliotransmitters can be detected by the generation of slow currents in neurons after the application of a studied drug in the presence of a blocker of voltage-gated sodium channels (VGSC), tetrodotoxin (TTX). In neurons, TTX suppresses spontaneous activity, which is manifested as the generation of action potentials. To discriminate the slow currents mediated by ionotropic glutamate receptors (iGluRs) and GABA_A_Rs, the membrane potential of the patched neuron is held at –30 mV to reverse the Cl^-^ gradient. If the Cl^-^ gradient in neurons is inverse, the activation of GABA_A_Rs causes Cl^-^ outflow from cells, which is registered as an outward current. In turn, activation of iGluRs induces an inflow of Na^+^ and Ca^2+^ that are registered as inward currents. Hence, Na^+^/Ca^2+^ and Cl^-^ flows under these conditions are referred to as slow inward (SIC) and slow outward (SOC) currents, respectively ^19,20^.

As our experiments demonstrated, the application of N^6^-CHA elicited a significant positive shift in the holding current and the generation of SOCs (Fig. 4A, A ′, B). The depolarization caused by Cl^-^ efflux from a neuron can explain these changes in the holding current. Application of a GABA_A_R antagonist, bicuculline, restored the holding current to the basal level, indicating that the observed shift was mediated by the activation of GABA_A_Rs. Since the spontaneous activity in neurons is suppressed with TTX under these conditions, GABA_A_Rs are activated by GABA released from astrocytes. Since GABA secretion by astrocytes is hypothesized to be linked to the activation of the PLC–PKC signaling pathway, we performed similar experiments in the presence of inhibitors targeting these proteins. The results showed that, in the presence of U73122 and Gö 6976, the application of the A_1_R agonist did not produce any significant changes in the holding current (Fig. 4C, D, E). This fact confirms our assumption about the impact of A_1_R agonists on the GABA release by astrocytes and the involvement of these cells in the realization of the antiepileptiform activity of A_1_R agonists.

**Figure 4.**
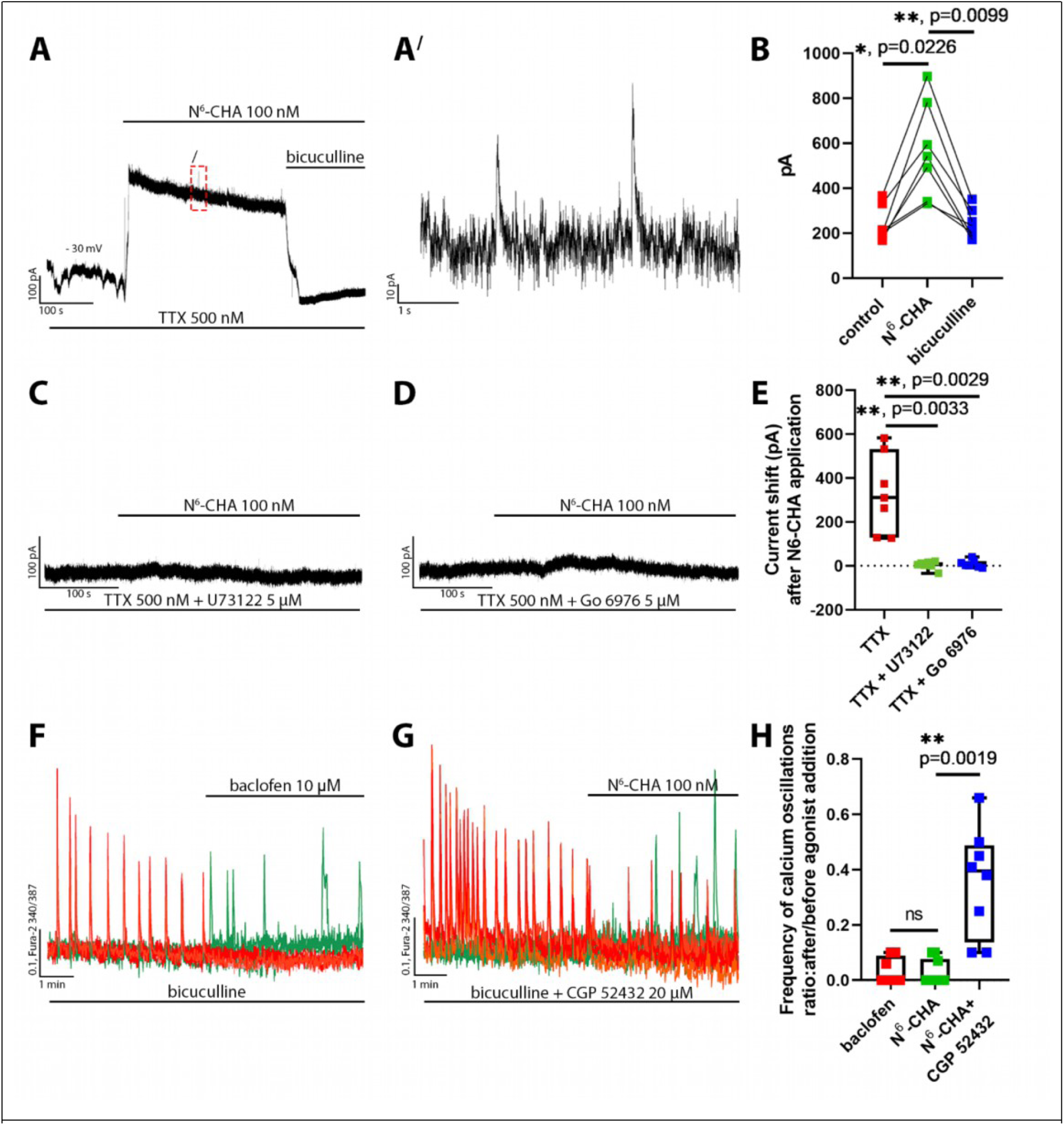
GABA release by astrocytes upon activation of A_1_Rs. (*A*) The effect of A_1_R agonist, N^6^-CHA (100 nM), and GABA_A_R antagonist on the holding current in a neuron in the presence of the VGSC blocker, TTX (500 nM). The membrane potential in a neuron was held at –30 mV. (***A***′) A magnified fragment of the electrophysiological recording marked with a red rectangle in Fig. 4A demonstrates representative SOCs. (***В***) The diagram shows the holding current values in the presence of N^6^-CHA and after bicuculline application. Friedman test followed by Dunn’s multiple comparisons test. The dots in the panel show the values of the holding current in individual experiments. ***(С, D)*** The effect of A_1_R agonist, N^6^-CHA (100 nM), on the holding current in a neuron in the presence of VGSC blocker, TTX (500 nM), and inhibitors of PLC (U73122, 5 μM) and PKC (Gö 6976, 5 μM), respectively. The membrane potential in a neuron was held at –30 mV. ***(E)*** Diagram showing current shift values after N^6^-CHA application in the presence of various inhibitors. The Kruskal‒Wallis test followed by Dunn’s multiple comparisons test. The dots in the panel show the values of the holding current in individual experiments. (***F***) The effect of GABA_B_R agonist, baclofen (10 μM), on the bicuculline-induced calcium oscillations. (***G***) The effect of GABA_B_R antagonist, CGP 52432 (20 μM), on the inhibitory action of the A_1_R agonist, N^6^-CHA. In panels C and D, red curves correspond to the responses of representative neurons, while green curves correspond to representative astrocytes from the same experiment. We analyzed 100 randomly chosen neurons and 100 astrocytes in each experiment. (***H***) The diagram shows the effects of the GABA(B)R agonist and antagonist on the frequency of calcium oscillations. The Kruskal‒Wallis test followed by Dunn’s multiple comparisons test. Dots on the diagram correspond to mean frequency values in individual experiments.

The blockade of GABA_A_Rs induced epileptiform activity in our experiments. In this case, GABA released by astrocytes can suppress neuronal activity only by acting on metabotropic GABA_B_ receptors (GABA_B_Rs). As Figure 4C shows, activation of GABA_B_Rs in the presence of bicuculline suppresses the calcium oscillations in neurons. Moreover, N^6^-CHA only slightly decreased the frequency of the bicuculline-induced calcium oscillations in the presence of GABA_B_R antagonist, CGP 52432 (20 μM) (Fig. 4D, E). Thus, these experiments also confirm the assumption about the contribution of astrocytic GABA release to the inhibitory action of A_1_R agonists.

### 3.3. The antiepileptic action of positive allosteric modulators of A_1_Rs

Our experiments showed that agonists of A_1_Rs have potential antiepileptic and neuroprotective activity. However, the use of orthosteric agonists *in vivo* is accompanied by various side effects. As considered, positive allosteric modulators (PAM) can minimize these effects. We have tested two commercially available PAMs – PD81723 and VCP171. As shown in Figure 5A, B, both A_1_R agonists (CCPA and N^6^-CHA) at the concentration 1 nM insignificantly affect the frequency of the bicuculline-induced calcium oscillations in neurons, while in the presence of the A_1_R PAM, the agonists almost completely suppressed the oscillations at this concentration (Fig. 5A, A′, B, B′). Moreover, our experiments demonstrated that PAMs significantly decrease the frequency of calcium oscillations, even in the absence of agonists (Fig. 5C, C′). Experiments with the A_1_R antagonist PSB36 confirmed that the observed effects of PAMs are mediated by A_1_Rs (Fig. 5D, D′).

**Figure 5.**
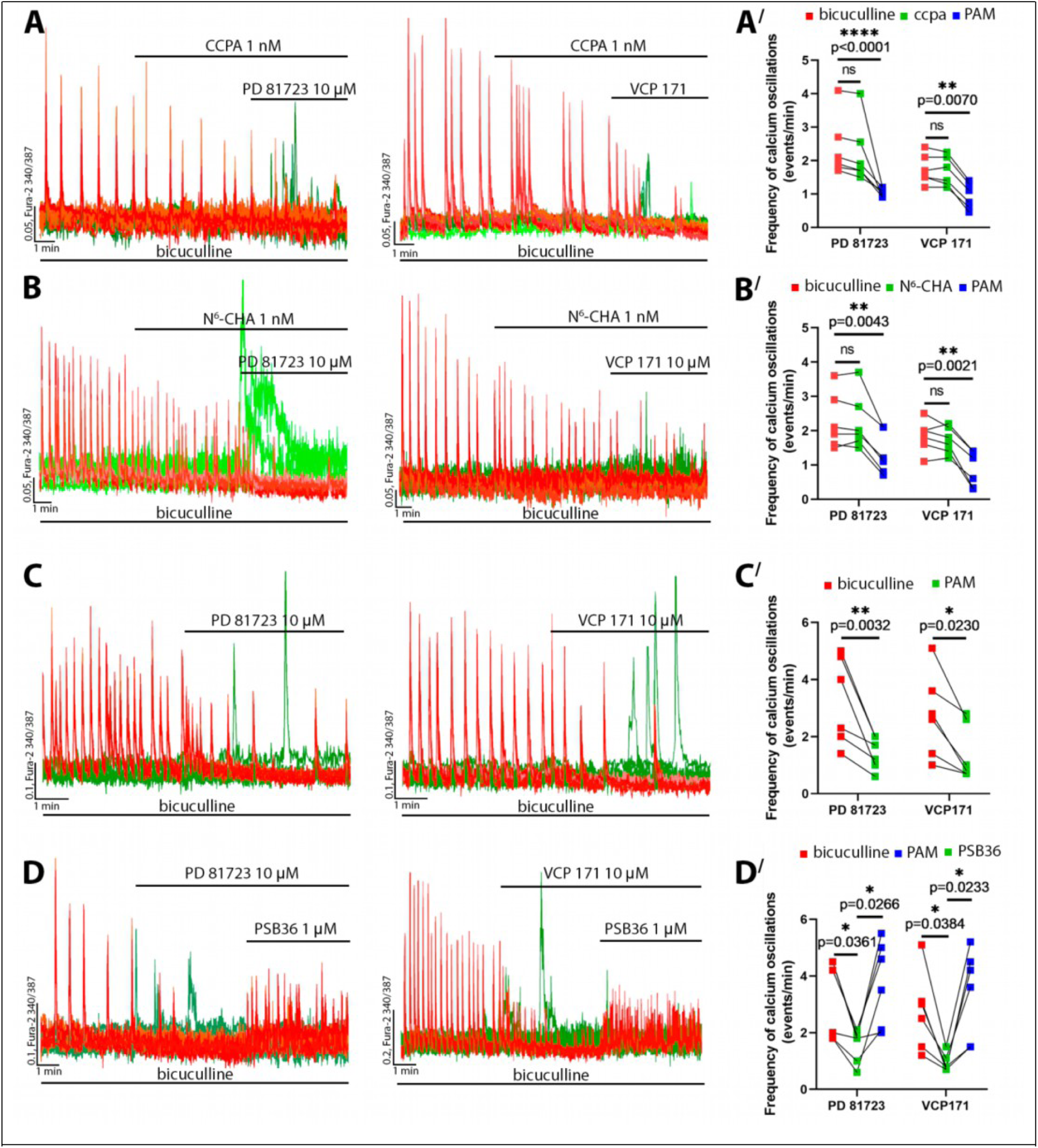
The effects of A_1_R positive allosteric modulators on the induced epileptiform activity. (***A***, ***В***) The effects of (***A***) CCPA (1 nM) and (***B***) N^6^-CHA (1 nM) on the bicuculline-induced (10 µM) calcium oscillations in neurons in the absence and presence of PD81723 (10 µM) and VCP171 (10 µM). (***A***′, ***В***′) Diagrams demonstrating the frequency of bicuculline oscillations in the presence of (***A***′) CCPA and (***В***′) N^6^-CHA before and after PD81723 or VCP171 application. Two-way ANOVA followed by Sidak’s multiple comparison test; (***С***) The effects of the A_1_R PAMs on the bicuculline-induced (10 µM) calcium oscillations in neurons in the absence of the A_1_R agonists. (***С***′) The frequency of the bicuculline-induced calcium oscillations in neurons before and after PD81723 (10 µM) and VCP171 (10 µM) application. Two-way ANOVA followed by Sidak’s multiple comparison test. (***D***) The effect of the A_1_R antagonist (PSB36, 1 µM) on the inhibitory action of PD81723 and VCP171 against bicuculline-induced calcium oscillations in neurons. Two-way ANOVA followed by Sidak’s multiple comparison test. In panels A, B, C, and D, red curves correspond to representative neurons, while green curves correspond to representative astrocytes. We analyzed 100 randomly chosen neurons and 100 astrocytes in each experiment. Dots on the diagram correspond to mean frequency values in individual experiments.

We have also tested newly synthesized and previously obtained substances with potential modulating activity on A_1_Rs. The detailed synthesis schemes are presented in the Supplementary section (see Supporting Information). It should be noted that substances from the SGA^21^ and TT ^22,23^ groups have been previously obtained; however, there is no information available regarding their biological effects on A_1_Rs. The key selection criterion was the similarity of the chemical structures of the studied substances to those of PD81723 and VCP171. As with commercially available A_1_R PAMs, the synthesized substances contain an aminothiophene scaffold. The substances with abbreviations ТТ8, ТТ28, and SGA396 (previously unknown compound) were the most effective suppressors of bicuculline-induced calcium oscillations (Fig. 6A, B, C).

**Figure 6.**
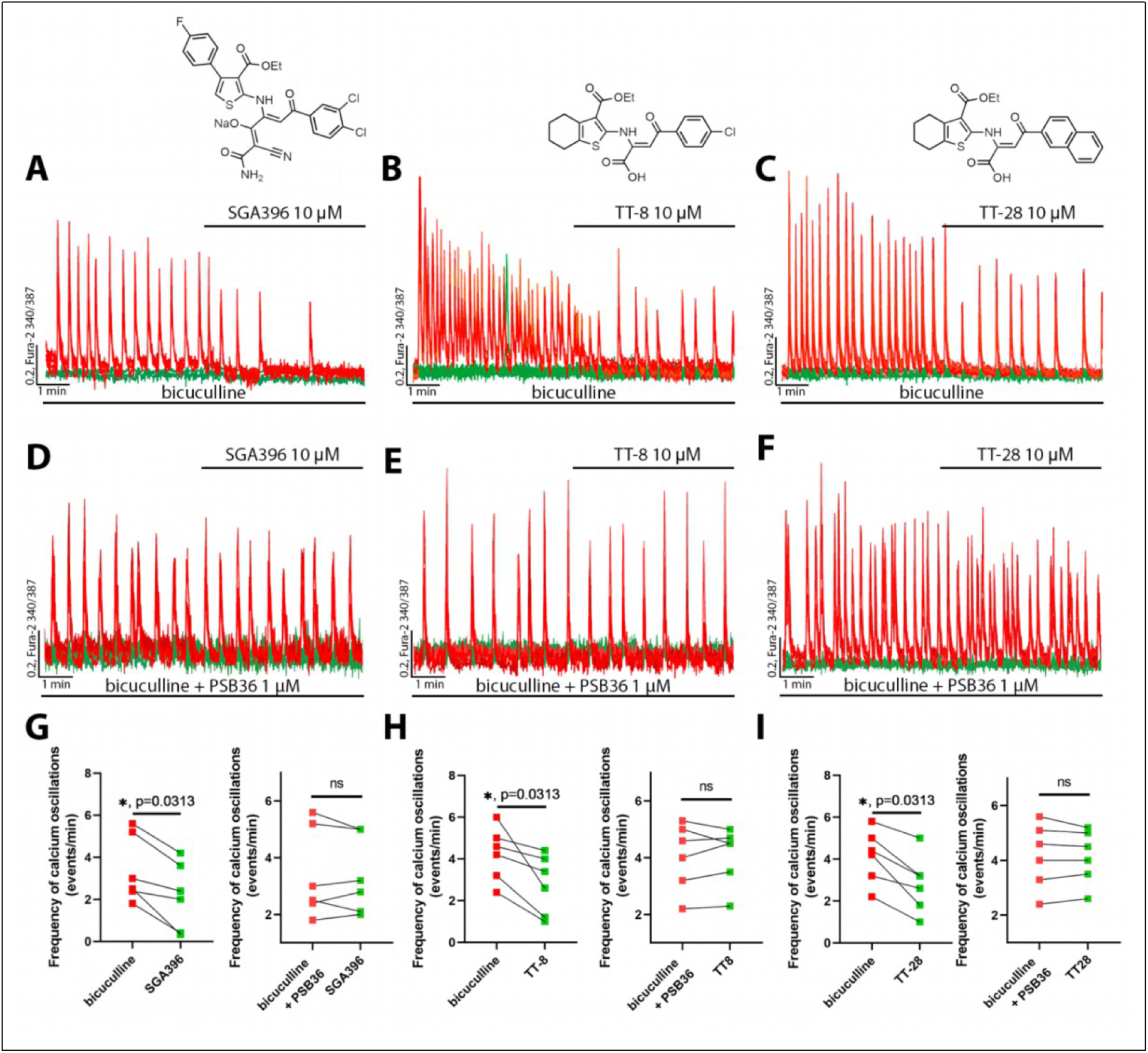
The effect of newly synthesized PAM of A1Rs. Panels *A*, *B*, and *C* demonstrate the effect of previously unknown PAMs on the bicuculline-induced calcium oscillations in neurons. The chemical structures of the PAMs are shown in the graphs above. (**G**, **H**, **I**) Diagrams illustrating the effects of the studied substances on the frequency of calcium oscillations in neurons in the absence and presence of the A1R antagonist PSB36. Wilcoxon matched-pairs signed rank test; 100 randomly chosen neurons and 100 astrocytes were analyzed. Red curves correspond to representative neurons, and green curves correspond to astrocytes. Dots in the diagrams correspond to the mean amplitude in individual experiments.

The selectivity of these compounds for A_1_ receptors was confirmed through inhibitory analysis (Fig. 6D–I). This effect was comparable to the action of PD81723 and VCP171. Thus, the obtained substances can be considered as potential neuroprotective drugs, whose efficacy will be further studied in *in vivo* research.

### 3.4. Evaluation of the antiepileptic activity of A_1_ receptor agonists and modulators *in vivo*

To evaluate the efficacy of A_1_R agonists under *in vivo* hyperexcitation conditions, we employed a seizure model induced by the systemic administration of picrotoxin (PTX), a GABA_A_R antagonist. Despite extensive literature on the effects of CCPA and N^6^-CHA in other models (e.g., kainate, pilocarpine), their impact in the context of disrupted GABAergic signaling has not been previously investigated. Our data demonstrate, for the first time, that both agonists – CCPA and N^6^-CHA – exert a strong, dose-dependent anticonvulsant effect, significantly improving survival in this model.

Analysis of the latency to the first seizure showed that even a single dose of N^6^-CHA at 0.5 mg/kg led to a noticeable delay in seizure onset: the median increased from 14 minutes in the control group to 48 minutes (Fig. 7B). At doses of 1 and 3 mg/kg, seizures were prevented in the majority of animals. CCPA exhibited a similar pattern, but required higher doses: at 1 mg/kg, half of the mice remained seizure-free during the entire observation period, and at 10 mg/kg, no seizures were observed at all (Fig. 7A). These differences were statistically significant (log-rank test, p < 0.0001). Estimates of hazard ratios (HRs) for time to seizure further supported the strong protective effects of both compounds. For N^6^-CHA, HRs remained consistently low across all doses (HR ≈ 0.02; 95% CI: 0.01–0.02), with complete within-group consistency (I² = 0%) (Fig. 7D). In contrast, CCPA showed slightly greater variability between doses (HR ranging from 0.54 to 0.02; I² = 70.7%), suggesting a less predictable dose–response relationship (Fig. 7C). Nevertheless, at the highest dose, its effect was comparable to that of N^6^-CHA.

**Figure 7.**
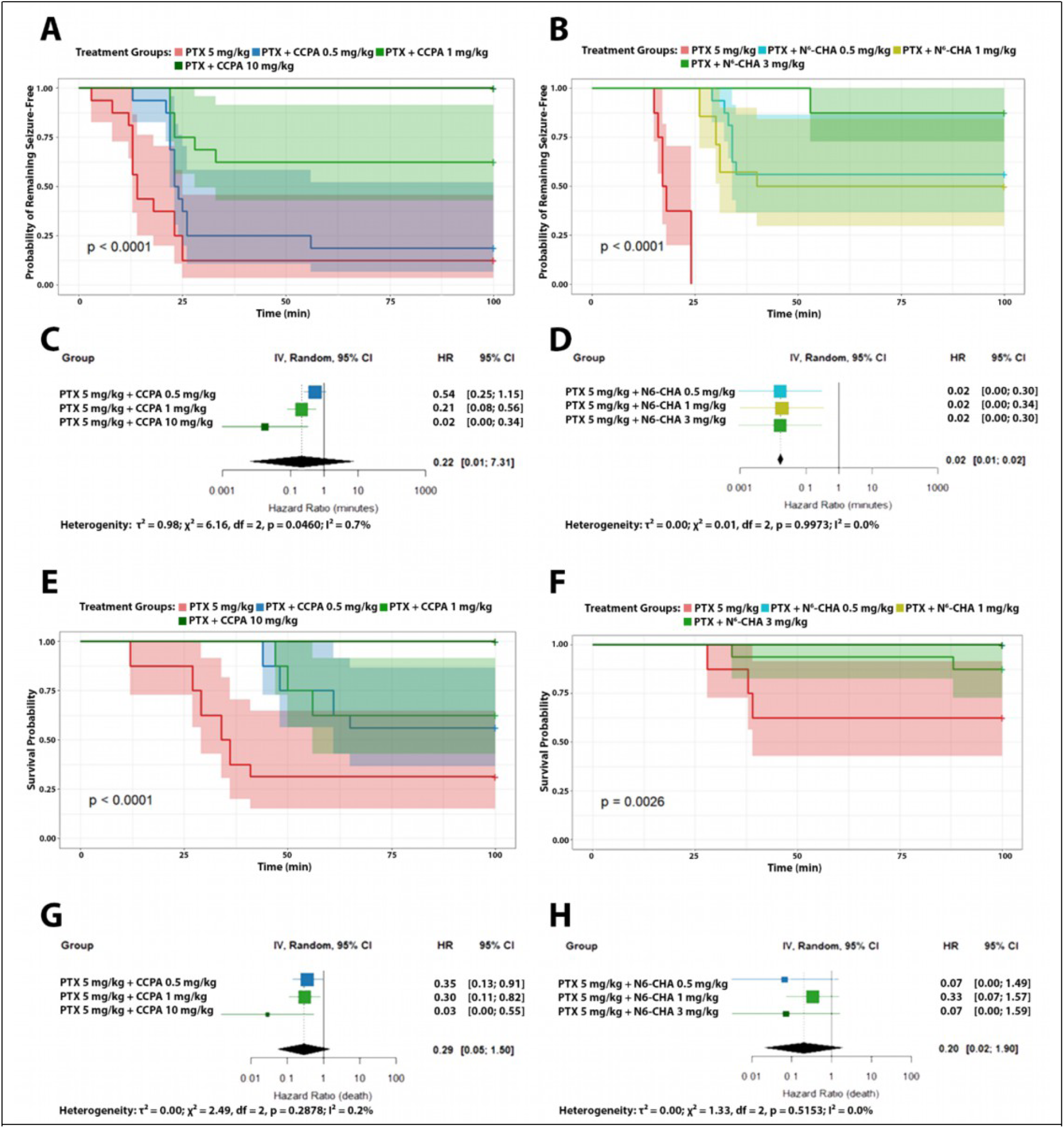
Effects of A_1_R agonists on seizure onset and survival in mice following picrotoxin injection. (***A***, ***B***) Kaplan–Meier curves representing the time to first seizure onset in mice pre-treated with various doses of A_1_R agonists (CCPA or N^6^-CHA) before picrotoxin (PTX) administration. Both CCPA (A) and N^6^-CHA (B) significantly delayed seizure onset in a dose-dependent manner. Notably, at higher doses (CCPA 10 mg/kg; N^6^-CHA 3 mg/kg), seizures were entirely prevented in most animals. (***C***, ***D***) – Forest plots showing hazard ratios (HRs) and 95% confidence intervals for seizure onset. Lower HR values indicate more substantial protective effects of the treatments. (***E***, ***F***) Kaplan–Meier survival curves following PTX injection. A_1_R agonists increased survival in a dose-dependent fashion, with N^6^-CHA (F) being especially effective even at 0.5 mg/kg. (***G***, ***H***) Forest plots presenting the impact of CCPA (G) and N^6^-CHA (H) on survival. Both compounds significantly reduced mortality, particularly at higher doses. In the N^6^-CHA groups, broader confidence intervals reflect the smaller group size and lower event rate.

These findings were also reflected in the survival data. Kaplan–Meier curves revealed a clear separation between the control and treated groups: in the PTX group, survival dropped sharply during the first hour, whereas administration of N^6^-CHA – even at 0.5 mg/kg – markedly increased the likelihood of survival, and at 3 mg/kg mortality was minimal (Fig. 7F). CCPA showed a similar trend, especially at 10 mg/kg (Fig. 7E). Hazard ratios for survival also confirmed the robust protective effects of both compounds: for both CCPA and N^6^-CHA, HR values were well below 1, indicating a significant reduction in the risk of death (Fig. 7G, H). Notably, no intergroup heterogeneity was observed (I² ≈ 0%), highlighting the consistency of the effects across animal populations. Altogether, our data provide the first evidence that activation of A_1_Rs can effectively counteract seizures and seizure-related lethality in a model based on impaired inhibitory transmission. N^6^-CHA appears particularly promising, as it exerts a reliable protective effect even at low doses and demonstrates high consistency across individuals.

These experiments were conducted using specialized cages equipped with both a video recording system and seismoacoustic sensors, allowing for the simultaneous monitoring of spontaneous locomotor activity (SLA) in mice. Figure 8 presents SLA recordings before and after seizure induction. Figure 8A shows the recording of a mouse that died following PTX administration, as indicated by the cessation of signal detection. In contrast, Figure 8A′ shows the recording of a mouse that survived the seizure; as the effect of PTX wore off, the animal’s normal circadian rhythm gradually resumed, characterized by alternating phases of rest and activity. A similar but less pronounced pattern was observed in mice pre-treated with N^6^-CHA at a dose of 0.5 mg/kg before PTX injection (Fig. 8B). However, when the N^6^-CHA dose was increased to 1 mg/kg, a marked disruption of locomotor rhythm was observed: the animals displayed either low activity levels or remained in a state of complete rest (Fig. 8C).

**Figure 8.**
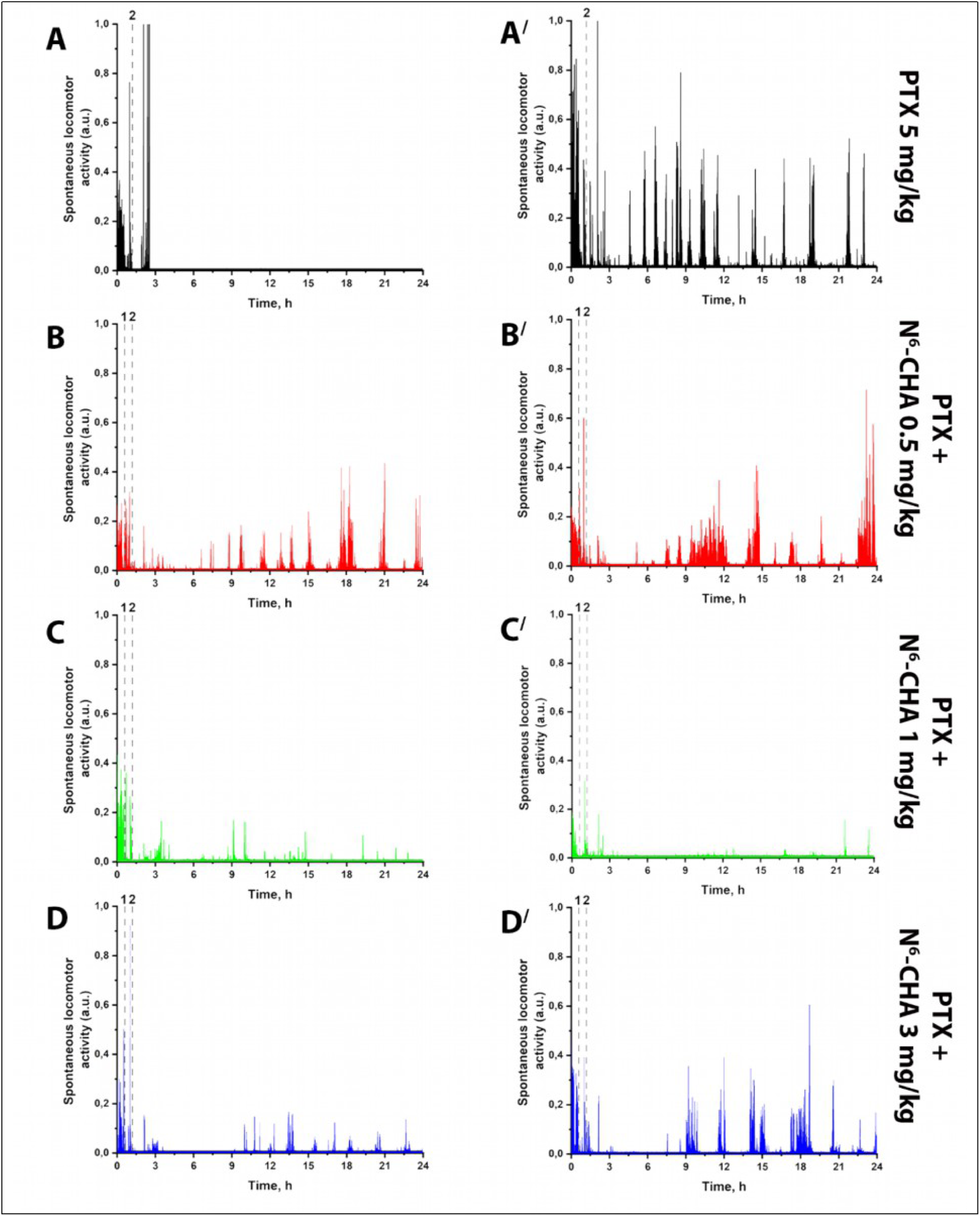
Dynamics of spontaneous locomotor activity (SLA) in mice before and after seizure induction. The figure shows SLA recordings from two mice in each group: (***A***, ***A′***) PTX 5 mg/kg; (B, B′) – PTX 5 mg/kg + N^6^-CHA 0.5 mg/kg; (C, C′) PTX 5 mg/kg + N^6^-CHA 1 mg/kg; (D, D′) PTX 5 mg/kg + N^6^-CHA 3 mg/kg. The dashed line labeled “1” indicates the time of N^6^-CHA injection, and the dashed line labeled “2” indicates the time of PTX injection.

By analyzing the SLA recordings and video data, we conditionally divided mouse activity into three states—low, moderate, and high activity (Fig. 9A). In the low activity state, mice are either sleeping or awake but immobile (threshold: 0–0.05 a.u.). Moderate activity includes behaviors such as grooming or eating without locomotion (threshold: 0.05–0.095 a.u.). High activity encompasses all movement-related behaviors within the cage, such as running, jumping, or digging (threshold: 0.095–1 a.u.). Based on this classification, we observed that mice treated with the A_1_R agonist mainly remained immobile during the first hour following the injection, showing significantly lower activity compared to control animals (Fig. 9B). Further analysis of post-PTX activity revealed characteristic differences between the groups. The graphs in Fig. 9C show the averaged SLA data for each group. Each point represents the mean activity level of the group within a 3-hour interval, with time counting starting 100 minutes after PTX injection. The data indicate that mice receiving the agonist at a dose of 1 mg/kg spent the majority of the first 24 hours in a low activity state (Fig. 9C), with this effect being most pronounced during the first 6 hours (Fig. 9D). These mice also exhibited reduced moderate and high activity levels, suggesting that A_1_R agonists have side effects that suppress overall locomotor activity. Similarly, mice that received the agonist at a dose of 3 mg/kg were mainly inactive or asleep during the first 6 hours post-injection. However, after this period, their SLA levels began to approach those of control animals.

**Figure 9.**
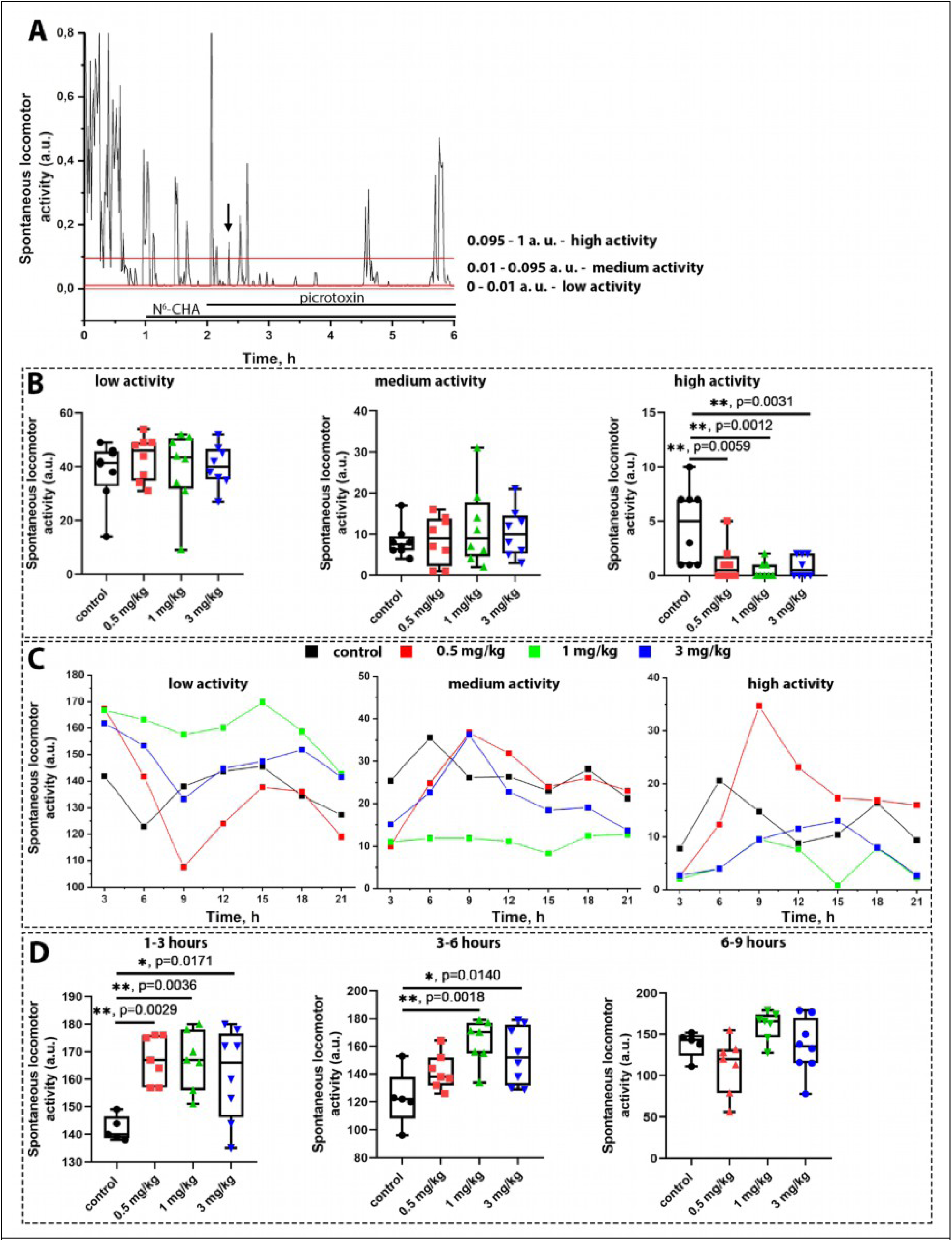
Analysis of SLA dynamics in mice following seizure induction. (***A***) Six-hour SLA recording illustrating threshold values used to categorize activity into three levels. An arrow indicates the moment of seizure onset. Red lines represent the thresholds (0.05 and 0.095 a.u.) separating low (0–0.05), moderate (0.05–0.095), and high (0.095–1) activity. (***B***) Comparative analysis of activity type distribution across groups during the first hour after PTX injection. Statistical analysis: ordinary one-way ANOVA followed by Dunnett’s multiple comparison test. (***C***) The temporal dynamics of SLA are presented as the average values of the three activity levels for each experimental group. (***D***) Comparative analysis of different activity types of defined time intervals following PTX administration. Statistical analysis: ordinary one-way ANOVA followed by Dunnett’s multiple comparison test.

To evaluate the anticonvulsant potential of A_1_R PAMs, we tested PD81723 and VCP171 in the PTX-induced seizure model. Unlike orthosteric agonists, PAMs do not directly activate the receptor; instead, they enhance the action of endogenous adenosine, potentially minimizing side effects. Single administration of either PD81723 (3 mg/kg) or VCP171 (10 mg/kg) did not significantly affect seizure latency or survival. As shown in Figure 10A–D, Kaplan–Meier analysis revealed no significant difference between treated and control animals (log-rank test: p = 0.43 for PD81723, p = 0.68 for VCP171 in seizure latency; p = 0.73 and p = 0.32, respectively, in survival). These data suggest that a single dose of PAMs may be insufficient to achieve a therapeutic effect in this model, potentially due to poor bioavailability or pharmacokinetic limitations.

**Figure 10.**
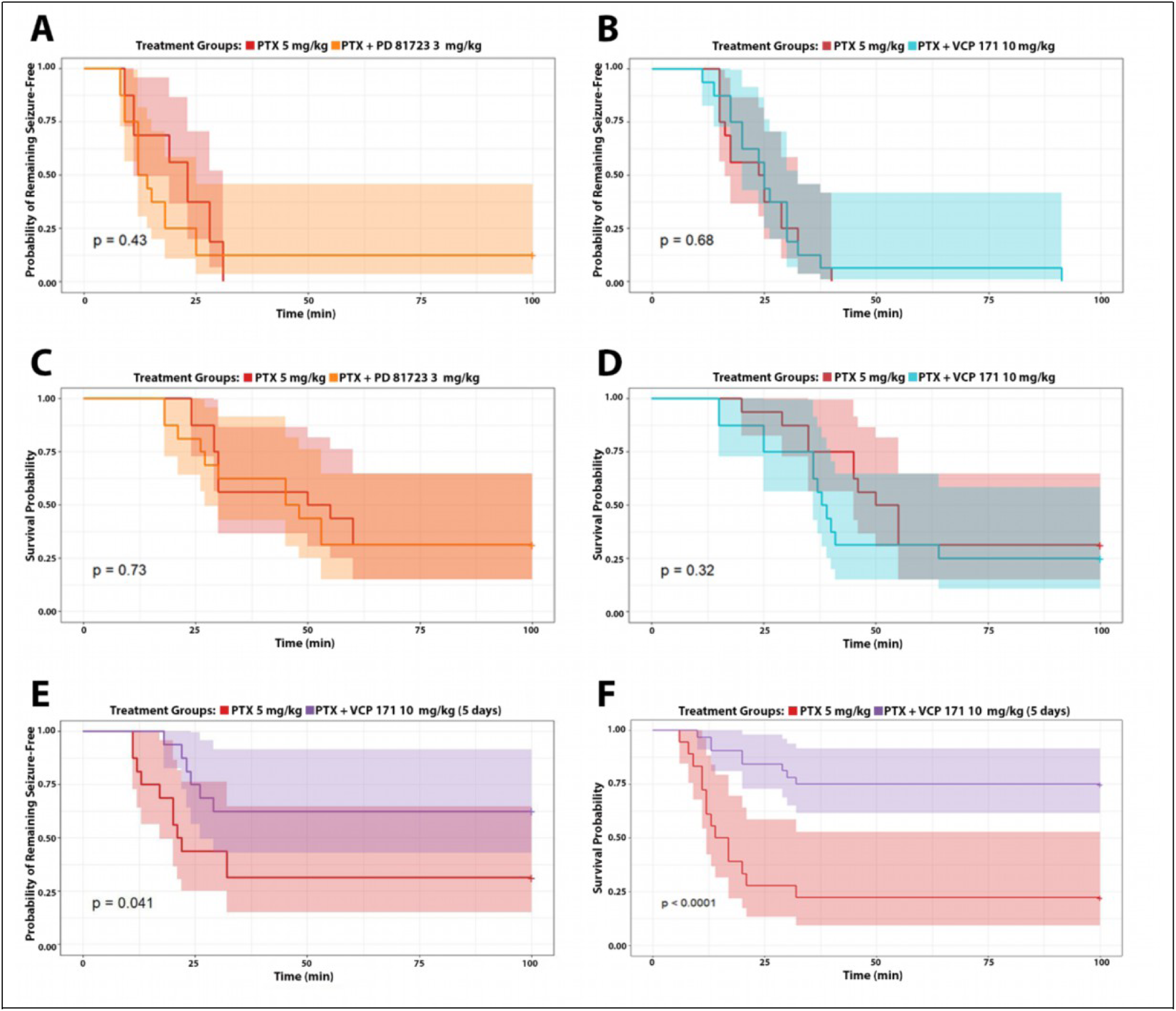
Effects of A1R positive allosteric modulators on seizure activity and survival in mice after picrotoxin administration. (***A***, ***B***) Kaplan–Meier curves showing the effects of single-dose administration of PD81723 (3 mg/kg) and VCP171 (10 mg/kg) on seizure latency. No significant differences were observed between the treated and control groups (log-rank test: p = 0.43 and p = 0.68, respectively). (***C***, ***D***) Survival probability after a single administration of the same compounds. Differences between groups did not reach statistical significance (p = 0.73 and p = 0.32). (***E***, ***F***) Effects of repeated VCP171 administration (10 mg/kg/day, 5 days, subcutaneously in oil): a significant delay in seizure onset (p = 0.041) and a marked increase in survival probability (p < 0.0001). Shaded areas represent 95% confidence intervals; “+” symbols indicate censored observations.

To address this, we tested a modified protocol involving repeated administration of VCP171 (10 mg/kg injection per day for 5 days, s.c. in oil). Under this scheme, a significant delay in seizure onset was observed (Figure 10E, p = 0.041), accompanied by a marked increase in survival probability (Figure 10F, p < 0.0001). These results indicate that prolonged exposure to VCP171 enables it to reach effective brain concentrations, allowing realization of its anticonvulsant potential. Altogether, while single administration of A_1_R PAMs was ineffective in this model, repeated dosing of VCP171 demonstrated significant protective effects, suggesting that pharmacokinetic optimization may be key for their *in vivo* efficacy.

To provide a comprehensive assessment of the efficacy of various A_1_R agonists and positive allosteric modulators (PAMs), we conducted a direct comparison of the most active compounds within a single experiment. The analysis included CCPA (10 mg/kg), N^6^-CHA (3 mg/kg), PD81723 (3 mg/kg), and VCP171 administered either as a single dose (10 mg/kg) or repeatedly over 5 days (10 mg/kg per day). Kaplan–Meier analysis revealed that all three active compounds — CCPA, N^6^-CHA, and VCP171 (5-day protocol) — significantly increased the latency to seizures (Figure 11A) and improved survival (Figure 11B) compared to the control group (PTX 5 mg/kg). In contrast, PD81723 and single-dose VCP171 did not exhibit significant protective effects.

**Figure 11.**
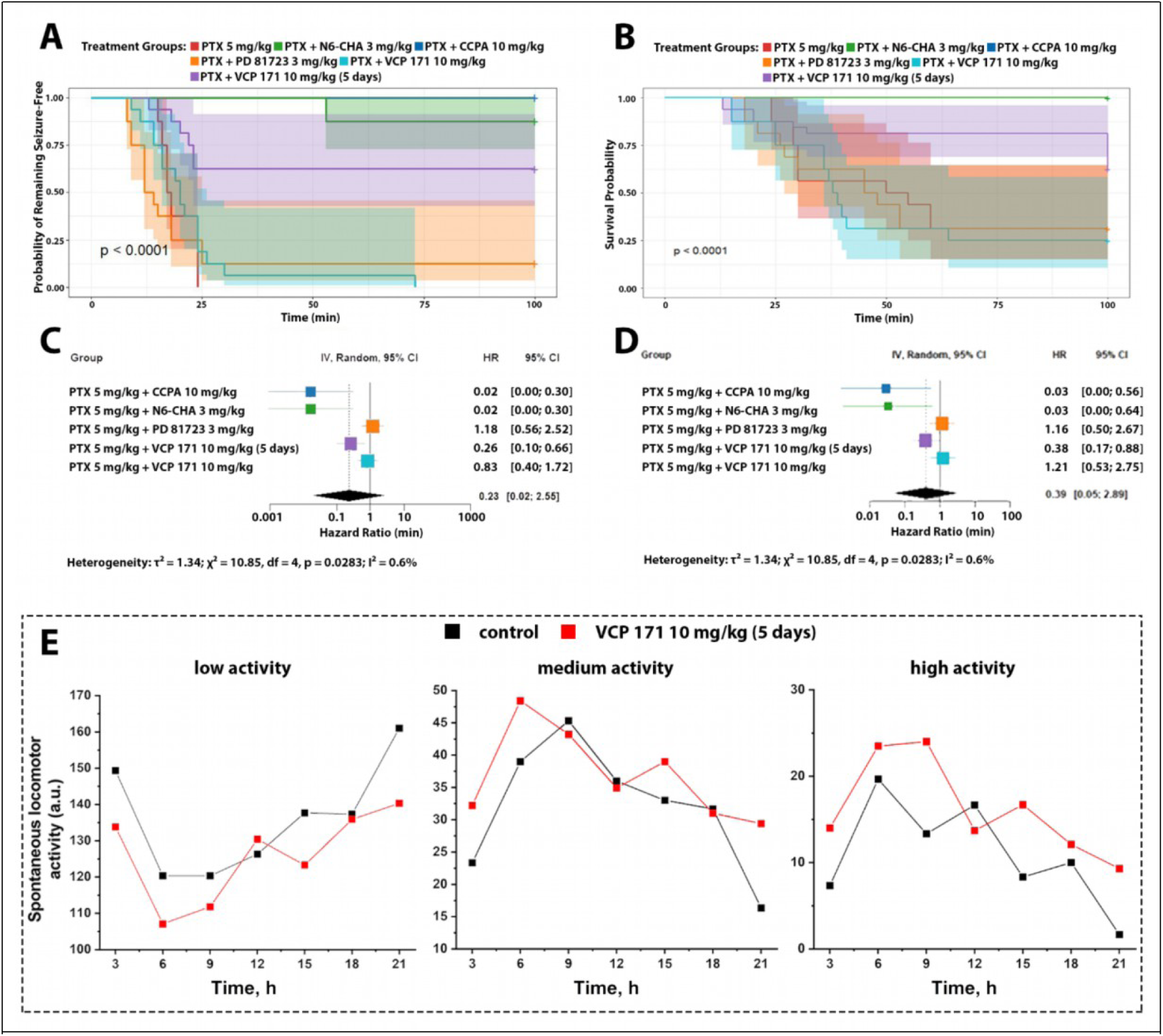
Comparative analysis of A_1_ receptor agonists and positive allosteric modulators in a picrotoxin-induced seizure model. (A, B) Kaplan–Meier survival curves showing the effects of various A_1_R ligands on seizure latency (A) and overall survival (B). CCPA (10 mg/kg), N^6^-CHA (3 mg/kg), and VCP171 (10 mg/kg, 5-day regimen) significantly delayed seizure onset and improved survival compared to the PTX-only group (log-rank test: p < 0.0001). In contrast, PD81723 and single-dose VCP171 did not show significant effects. (C, D) Forest plots displaying hazard ratios (HRs) and 95% confidence intervals for seizure latency (C) and survival (D). The most potent protection was observed with N^6^-CHA and CCPA (HR ≈ 0.02). Repeated VCP171 administration also produced a strong effect (HR 0.26 and 0.38 for seizure and survival, respectively). Between-group heterogeneity was low (I² < 1%). (E) Spontaneous locomotor activity (SLA) analysis over 21 hours post-PTX injection. Activity was categorized as low, medium, or high. Mice treated with VCP171 (5 days) showed preserved circadian motor rhythms and reduced behavioral suppression compared to controls, particularly in the high-activity phase.

To quantitatively evaluate the treatment outcomes, hazard ratios (HRs) were calculated for each compound. As shown in Figures 11C–D, the most pronounced reduction in seizure and mortality risk was observed for N^6^-CHA and CCPA, both with HRs close to 0.02. Repeated administration of VCP171 also conferred a robust protective effect (HR for seizures: 0.26; for survival: 0.38), comparable to that of the agonists. Single-dose PD81723 and VCP171 did not affect survival outcomes (HR ≈ 1). Intergroup heterogeneity was minimal (I² < 1%), indicating consistent effects across experimental groups.

In addition, spontaneous locomotor activity (SLA) was monitored over 21 hours following PTX administration (Figure 11E). In the control group, a sharp drop in activity was observed shortly after injection, particularly in the high-activity category, reflecting seizure development and subsequent behavioral suppression. In contrast, mice that received VCP171 for 5 days maintained more stable activity levels in the medium and high ranges, closely resembling the physiological circadian rhythm. These results suggest that the protective effects of VCP171 are achieved without the marked motor suppression observed with high doses of the A_1_R agonists.

Altogether, these findings demonstrate that both orthosteric agonists and positive allosteric modulators of A_1_Rs can effectively prevent seizure development and reduce mortality when administered under an optimized regimen. Notably, repeated administration of VCP171 combines therapeutic efficacy with a more favorable behavioral profile, supporting its potential as a promising candidate for further investigation.

## 4. Discussion

### 4.1. Mechanism of the inhibitory action of A_1_R agonists

In this work, it was demonstrated using three different models of hyperexcitation that A_1_R agonists can have both antiepileptic and neuroprotective effects, including the prevention of calcium overload in specific neuronal populations. In the case of bicuculline-induced epileptiform activity, hyperexcitation emerges from disinhibition of the neuronal network due to the blockade of GABA_A_Rs. Our previous study shows that calcium oscillations under these conditions are caused by a paroxysmal depolarization shift resulting primarily from the activation of AMPA receptors ^6^. Consequently, the oscillations occur due to increased activity of glutamatergic neurons, which become hyperactivated in the absence of GABA_A_R-mediated inhibition. Such abnormal activity in neuron-glial cultures is considered a cellular correlate of interictal spikes recorded on EEG during epilepsy development or between seizures ^24^. We have previously shown that A_1_R agonists can both suppress and prevent the occurrence of calcium oscillations ^6^, and in this work, we elucidate the mechanisms underlying this inhibitory action. Our experiments suggest that the suppression of epileptiform activity is achieved not only through the activation of neuronal but also astrocytic A_1_Rs.

In the hippocampus, including hippocampal cultures, the expression of A_1_Rs has been demonstrated in both neurons and astrocytes ^25^. In addition to inhibiting adenylyl cyclase, leading to a decrease in cAMP levels, activation of A_1_Rs in neurons causes a negative modulation of VGCCs and activation of GIRK channels ^26^. Using acute slices and cultures of hippocampal neurons, it was demonstrated that activation of A_1_Rs leads to a decrease in the frequency of miniature spontaneous synaptic currents (mPSCs) ^27,28^. This effect is calcium-independent and is not associated with cAMP levels or PLC activity. Presumably, the decrease in mPSC frequency is caused by the direct interaction of the Gβγ subunit with SNARE proteins responsible for vesicle fusion with the presynaptic membrane ^26^. Activation of A_1_Rs can also lead to positive modulation of potassium K_ATP_ and SK channels, as demonstrated in substantia nigra neurons and retinal ganglion cells, respectively ^29,30^. However, it is not known whether a similar change in the activity of these potassium channels occurs in hippocampal neurons.

Despite the numerous potential effects on neurons, our experiments suggest that astrocytes play a crucial role in the inhibitory effect of A_1_R agonists on induced hyperexcitation. This conclusion is based on several facts: firstly, the addition of N^6^-CHA enhances calcium activity in astrocytes, increasing both the number of astrocytes generating the calcium transients and the frequency of these events; secondly, the inhibitor of the Gβγ subunit, gallein, reduce the effectiveness of A_1_Rs agonists, while no change in calcium activity in astrocytes is observed; thirdly, N^6^-CHA application in the presence of TTX blocking neuronal spontaneous firing activity causes a GABA_A_Rs-mediated positive shift in the holding current and SOCs occurrence in neurons; fourthly, the inhibitory activity of the A_1_R agonist against induced epileptiform activity is attenuated in the presence of GABA_B_R antagonist. Based on these data, we hypothesize that activation of A_1_Rs triggers the Gβγ–PLC signaling pathway and mobilization of Ca^2+^ from the ER. In turn, the A_1_R-mediated [Ca^2+^]_i_ elevations in astrocytes stimulate the release of GABA, which contributes to the suppression of epileptiform activity in neurons. We have previously demonstrated a similar mechanism of gliotransmission upon activation of α_2_-adrenergic receptors ^14,18^. Several studies have shown a correlation between increased [Ca^2+^]_i_ in astrocytes and their release of various mediators; however, the intermediate steps between these two events have not been established ^31^. We have shown that inhibition of PKC significantly reduces the antiepileptiform activity of A_1_R agonists but does not affect the calcium activity of astrocytes. This may indicate that GABA release by astrocytes is a PKC-dependent process: activation of astrocytic A_1_Rs evokes a [Ca^2+^]_i_ increase, but GABA release does not occur, as it apparently requires PKC activation. Since no studies demonstrate the involvement of PKC in gliotransmission, the exact mechanism of this process requires further investigation.

The inhibitory effect of A_1_R agonists was also demonstrated in a model of hyperexcitation induced by NH_4_Cl. As we have previously shown, calcium oscillations in this case arise due to the depolarization of neurons caused by ammonium ions ^18^. Unlike the bicuculline-induced calcium oscillations, the NH_4_Cl-induced oscillations have a higher frequency and are accompanied by an increase in the basal [Ca^2+^]_i_ level. Nevertheless, application of A_1_R agonists suppresses even such high-frequency activity and decreases [Ca^2+^]_i_ to baseline values. As with the bicuculline-induced calcium oscillations, the agonist enhanced the calcium activity of astrocytes. In this regard, it can be assumed that, as in the case of epileptiform activity induced by bicuculline, under conditions of hyperammonemia, the addition of the agonist leads to the activation of A_1_Rs on both neurons and astrocytes, and their combined activity results in the suppression of hyperexcitation.

Unlike the other two models of hyperexcitation, the A_1_R agonist did not affect the calcium activity in astrocytes upon acute glutamate excitotoxicity. Notably, glutamate evoked in astrocytes a response whose pattern is similar to the pattern of responses to activation of various metabotropic receptors (biphasic change in [Ca^2+^]_i_, consisting of a peak and plateau). Apparently, the response is mediated by the activation of metabotropic glutamate receptors (mGluRs) ^32^. Similarly to transients evoked by A_1_Rs activation, the internal stores of the endoplasmic reticulum act as the source of Ca^2+^ for such a response. This fact may explain the absence of the reaction in astrocytes when N^6^-CHA was added after glutamate washout. Nevertheless, activation of A_1_Rs accelerates the restoration of calcium homeostasis in neurons after acute glutamate excitotoxicity. Hence, activation of neuronal A_1_Rs mitigates dysregulation of calcium homeostasis even without an astrocytic response. Since glutamate continues to be secreted by hyperexcited neurons even after washout, activation of A_1_Rs may help suppress this process by reducing cAMP levels. In addition, the activation of various potassium channels, such as GIRK or SK, as well as the suppression of VGCC activity, may also contribute to the restoration of neuronal ion homeostasis through hyperpolarization and a reduction in Ca^2+^ influx ^26^. Since calcium overload can lead to cell death ^33^, accelerating the restoration of calcium homeostasis after glutamate toxicity can be considered a neuroprotective effect, which requires further detailed consideration beyond the scope of this study.

### 4.2. The effect of positive allosteric modulators of A_1_Rs

Considering our data and the results of other studies, it can be argued that A_1_Rs represent a promising target for the development of drugs that exert anticonvulsant or neuroprotective effects. Nevertheless, as with orthosteric agonists of other Gi-coupled receptors, such as cannabinoid receptors, GABA_B_Rs, and mGluRs, the use of A_1_R agonists *in vivo* is associated with various undesirable side effects. In the case of A_1_R agonists, these effects include bradycardia, hypotension, and alterations in renal hemodynamics ^34^. However, as studies demonstrate, side effects can be avoided through the use of positive allosteric modulators. For example, the A_1_R positive modulator TRR469 exhibited an antinociceptive effect without inducing catalepsy or side effects on motor activity, as observed with the direct agonists ^15^. In the case of PAMs of other G_i_-coupled receptors, a similar effect was observed: PAMs of cannabinoid and GABA_B_Rs exhibited an analgesic effect without pronounced side effects ^35–37^.

Regarding the action of PAMs in various pathologies associated with hyperexcitation, such as epilepsy, ischemia, and traumatic brain injury, only a single study devoted to this issue has been published. There is extensive evidence suggesting that activation of A_1_Rs has both anticonvulsant ^12^ and neuroprotective effects in cerebral ischemia ^38^. However, as mentioned above, clinical studies of A_1_R agonists were limited by side effects, primarily on the cardiovascular system. To date, only one study has demonstrated a neuroprotective effect of the A_1_R PAM, PD81723, in a model of cerebral ischemia ^39^. Our experiments demonstrate that the concentration of endogenous adenosine, which increases upon hyperexcitation, is sufficient for the realization of the antiepileptic action of PAMs. Various studies have shown that the extracellular concentration of adenosine increases in cerebral ischemia ^8,40^, epilepsy ^13^, and after traumatic brain injury ^41,42^. Thus, the use of PAMs can enhance the neuroprotective effect of endogenously produced adenosine, while having no pronounced effect on other organs where its concentration remains at physiological levels.

### 4.3. The effect of A_1_ receptor agonists and PAMs on picrotoxin-induced seizures

The anticonvulsant effects of various A_1_R agonists have previously been demonstrated in pilocarpine and kainate models of epilepsy ^43–45^. However, their efficacy in seizure models based on GABA_B_R inhibition has not been studied. In this work, we investigated the anticonvulsant properties of A_1_R agonists *in vivo* using picrotoxin, a specific GABA_A_R antagonist, which parallels our previously established *in vitro* model. Both tested A_1_R agonists demonstrated a robust anticonvulsant effect, as evidenced by a significant increase in the latency to seizure onset and improved survival rates. In some cases, seizures were prevented entirely, further supporting the therapeutic potential of A_1_Rs as targets for the development of antiepileptic drugs.

However, our findings also showed that the use of direct A_1_R agonists is associated with several side effects, particularly dose-dependent suppression of spontaneous locomotor activity (SLA), most prominent during the first 6 hours post-injection. Notably, at a dose of 0.5 mg/kg N^6^-CHA, which was sufficient to achieve an anticonvulsant effect, the reduction in activity was minimal and mainly reflected by an increase in rest or sleep time, with a return to normal circadian activity patterns within 3–6 hours.

To minimize side effects, we also evaluated PAMs of A_1_ receptors—PD81723 and VCP171. Due to their high lipophilicity, these compounds were administered in an oil-based vehicle, which may have limited their bioavailability. Single-dose administration of PAMs did not significantly affect seizure onset or survival. However, chronic administration of VCP171 over five consecutive days resulted in a marked increase in latency to seizure onset and a significant improvement in post-seizure survival. Notably, even after repeated administration, VCP171 did not induce any notable side effects, highlighting its promise as a candidate for further preclinical evaluation.

## Conclusion

Overall, our study elucidates the molecular mechanisms underlying the antiepileptic activity of A1R agonists, highlighting their complexity and revealing the pivotal role of neuron-glial interactions in this process. Moreover, our data indicate that A_1_R PAMs have great potential for the design of novel anticonvulsant and neuroprotective drugs. Nevertheless, further studies are required to confirm this assumption and identify the potential positive and negative effects of such compounds.

## Data availability statement

The data that support the findings of this study are available from the corresponding author upon reasonable request.

---

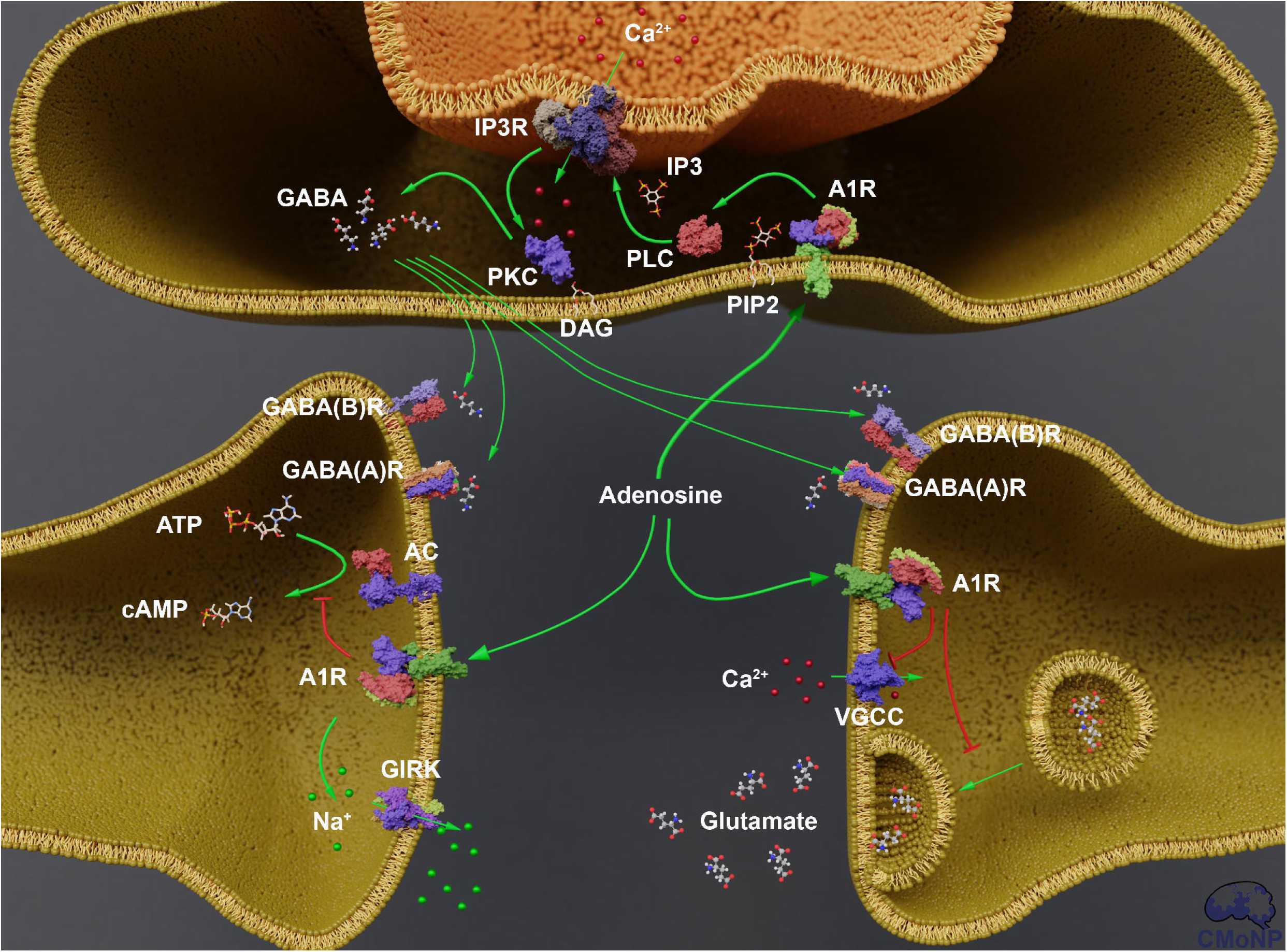

