## supplementary files for "Neuroprotective action of agonists and modulators of A_1_ adenosine receptors upon hyperexcitation: mechanism of the antiepileptic activity and role of neuron-glial interaction"

**1. Synthesis and characterization of compounds.**

All the air-sensitive reactions were carried out under nitrogen atmosphere in standard Schlenk flasks. All the chemicals purchased from Sigma-Aldrich and used without further purification. 4′-chloroacetophenone (97%), 3′,4′-dichloroacetophenone (99%), 2-acetonaphthone (99%), diethyl oxalate (≥99%), sodium methoxide (95%), cyclohexanone (≥99.0%), 4′‑fluoroacetophenone (99%), sulfur (99.98%), morpholine (≥99%), ethyl cyanoacetate (≥98%), DCC (99%), cyanoacetamide (99%), sodium carbonate (≥99.5%) were purchased from Sigma‑Aldrich. Methanol (MeOH), acetonitrile, ethanol (EtOH), and dioxane were dried over activated molecular sieves (3 Å) in Erlenmeyer flasks for two days prior to use. Chemically pure grade solvents were subjected to additional purification and drying. All other reagents were purchased from various commercial sources and used without additional purification.

NMR spectra were recorded on Bruker Avance III (400 MHz). The chemical shifts (δ) were measured in ppm with respect to the solvent (CDCl_3_, ^1^Н: δ = 7.26 ppm, ^13^C: δ = 77.2 ppm; DMSO‑*d_6_*, ^1^Н: δ = 2.50 ppm, ^13^C: δ = 39.5 ppm). Elemental analyses were performed with a Leco CHNS-932 elemental analyzer. Melting points were recorded with a Stuart SMP 40.

**2. Synthesis of compounds**

Starting (*Z*)-4-R^3^-2-hydroxy-4-oxobut-2-enoic acid **4a-с** are known compounds and were synthesized following the protocols from literature [S1].

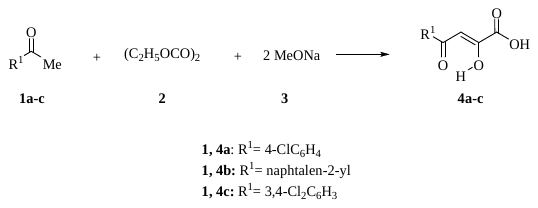

**Scheme S1.** Synthesis of (*Z*)-4-R^3^-2-hydroxy-4-oxobut-2-enoic acid **4a-с**.

Starting ethyl 2-amino-4,5,6,7-tetrahydrobenzo[*b*]thiophene-3-carboxylate **8a** is known compound and was synthesized following the protocols from literature [S2].

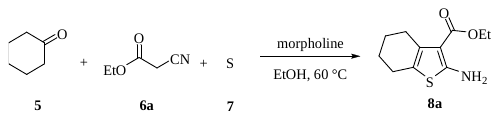

**Scheme S2.** Synthesis of ethyl 2-amino-4,5,6,7-tetrahydrobenzo[*b*]thiophene-3-carboxylate **8a**.

Starting ethyl 2-amino-4-(4-fluorophenyl)thiophene-3-carboxylate **8b** is known compound and was synthesized following the protocols from literature [S3].

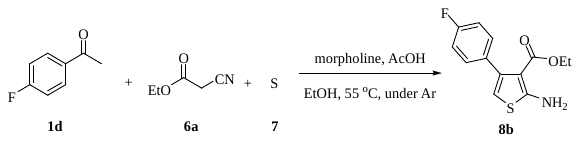

**Scheme S3.** Synthesis of ethyl 2-amino-4-(4-fluorophenyl)thiophene-3-carboxylate **8b**.

A solution of ethyl 2-amino-4,5,6,7-tetrahydrobenzo[*b*]thiophene-3-carboxylate **8a** (1 mmol) in 3 mL of ethanol was added to a solution of corresponding compounds **4a,b** (1 mmol) in 3 mL of ethanol. The resulting mixture was heated to 60 °C for 1 hour. After 1 hour the solution was cooled to –27 °C, and the precipitate was filtered off and recrystallized from acetonitrile providing compounds **9a,b**.

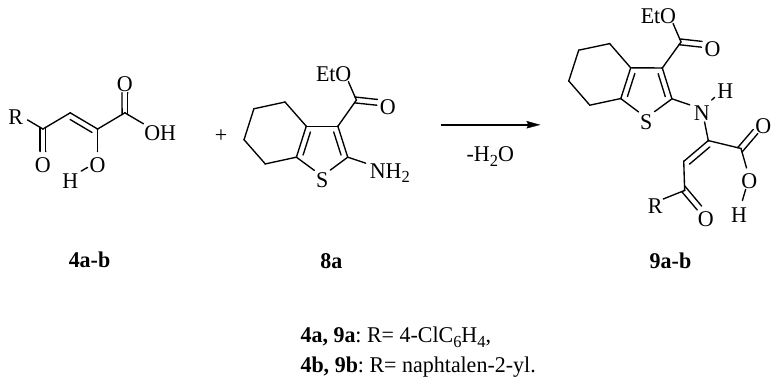

**Scheme S4.** Synthesis of compounds **9a,b**.

A mixture of compounds **4c** (1 mmol), **8b** (1 mmol) and DCC (2 mmol) was dissolved in 20 ml of anhydrous dioxane; the resulting solution was refluxed for 2 h and intensely stirred. After 2 h from the hot solution, precipitate urea was filtered. Then the solution was cooled to r.t. and treated with 2-cyanoacetamide (1 mmol), followed by addition of sodium carbonate (1 mmol). The solution was heated to 50 °C and maintained for 24 h. Then the solution was cooled to –27 °C, the precipitate was filtered off and recrystallized from acetonitrile providing compound **10**.

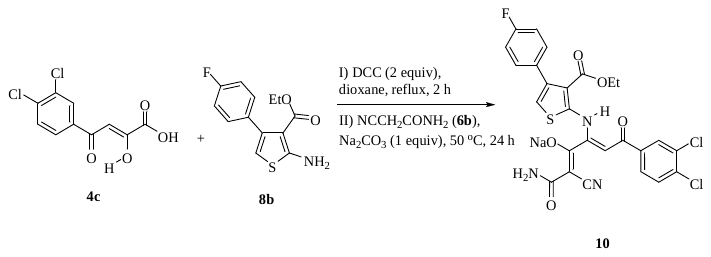

**Scheme S5.** Synthesis of sodium 1-amino-2-cyano-6-(3,4-dichlorophenyl)-4-((3-(ethoxycarbonyl)-4-(4-fluorophenyl)thiophen-2-yl)amino)-1,6-dioxohexa-2,4-dien-3-olate **10**.

**3. Compounds characterization**

*3.1. NMR spectra analysis.*

*(Z)-4-(4-chlorophenyl)-2-hydroxy-4-oxobut-2-enoic acid (****4a****)*. White crystals; 35.80 g, 79% yield; m.p. 163–164 °C. (lit. 163–165 °C [S4]) ^1^H NMR (CDCl_3_, 400 MHz) δ (ppm): 7.99 (m, 2H), 7.59 (m, 2H), 7.08 (s, 1H).

*(Z)-2-hydroxy-4-(naphthalen-2-yl)-4-oxobut-2-enoic acid (****4b****)*. White crystals; 20.34 g, 83% yield; m.p. 184–185 °C (lit. 184–186 °C [S5]). ^1^H NMR (DMSO-*d*_6_, 400 MHz) δ (ppm): 8.19 (m, 1H), 8.07 (m, 1H), 8.00 (m, 2H), 7.67 (m, 3H), 7.28 (s, 1H).

*(Z)-4-(3,4-dichlorophenyl)-2-hydroxy-4-oxobut-2-enoic acid (****4c****)*. White crystals; 36.78 g, 71% yield; m.p. 168–169 °C (lit. 170–171 °C [S6]). ^1^H NMR (DMSO-*d*_6_, 400 MHz) δ (ppm): 8.25 (m, 1H), 8.02 (m, 1H), 7.82 (m, 1H), 7.02 (s, 1H).

*Ethyl 2-amino-4,5,6,7-tetrahydrobenzo[b]thiophene-3-carboxylate (****8a****)*. Yellow crystals; 19.15 g, 85% yield; m.p. 112–113 °C. (lit. 111–115 °C [S7]). ^1^H NMR (CDCl_3_, 400 MHz) δ (ppm): 5.93 (s, 2H), 4.28 (q, *J* = 7.1 Hz, 2H), 2.73 (m, 2H), 2.52 (m, 2H), 1.79 (m, 4H), 1.36 (t, *J* = 7.1 Hz, 3H).

*Ethyl 2-amino-4-(4-fluorophenyl)thiophene-3-carboxylate (****8b****)*. Yellow crystals; 19.35 g, 73% yield; m.p. 92–93 °C. ^1^H NMR (CDCl_3_, 400 MHz) δ (ppm): 7.16 (m, 2H), 6.92 (m, 2H), 6.01 (br. s, 2H), 5.95 (s, 1H), 3.97 (q, *J* = 7.1 Hz, 2H), 0.89 (t, *J* = 7.1 Hz, 3H).

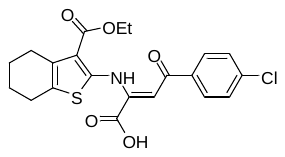

**ТТ8**

*4-(4-Chlorophenyl)-2-((3-(ethoxycarbonyl)-4,5,6,7-tetrahydrobenzo[b]thiophen-2-yl)amino)-4-oxobut-2-enoic acid (****9a****)*. Red solid (yield 90%), m.p. 176–177 °С (lit. 176–177 °С [S8]).
^1^H NMR (CDCl_3_, 400 MHz) δ (ppm): 12.33 (s, 1Н), 7.98 (m, 2H), 7.51 (m, 2H), 7.08 (s, 1H), 4.41 (q, *J* = 7.1 Hz, 2H), 2.84 (m, 2H), 2.78 (m, 2H), 1.85 (m, 4H), 1.41 (t, *J* = 7.1 Hz, 3H). ^13^C NMR (CDCl_3_, 100 MHz) δ (ppm): 188.1, 164.3, 162.0, 146.3, 143.8, 140.2, 135.9, 134.8, 129.9, 129.7, 129.3, 118.5, 96.1, 61.4, 26.5, 24.9, 22.7, 22.4, 14.2.

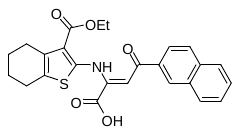

**ТТ28**

*2-((3-(ethoxycarbonyl)-4,5,6,7-tetrahydrobenzo[b]thiophen-2-yl)amino)-4-(naphthalen-2-yl)-4-oxobut-2-enoic acid (****9b****)*. Red solid (yield 89%), m.p. 116–117 °С. (lit. 116–118 °C [S9]). ^1^H NMR (CDCl_3_, 400 MHz) δ (ppm): 12.13 (s, 1H), 8.47 (m, 1H), 8.03 (m, 1H), 7.90 (m, 2H), 7.59 (m, 3H), 7.04 (s, 1H), 4.44 (q, *J* = 7.1 Hz, 2H), 2.84 (m, 2H), 2.65 (m, 2H), 1.81 (m, 4H), 1.43 (t, *J* = 7.1 Hz, 3Н). ^13^C NMR (CDCl_3_, 100 MHz) δ (ppm): 194.3, 164.3, 162.3, 146.0, 144.1, 137.7, 134.9, 134.1, 133.0, 130.3, 129.8, 128.8, 127.9, 127.8, 126.9, 125.8, 124.9, 118.8, 101.3, 61.5, 26.6, 25.0, 22.9, 22.6, 14.4.

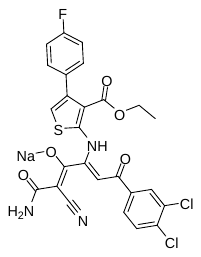

**SGA396**

*Sodium 1-amino-2-cyano-6-(3,4-dichlorophenyl)-4-((3-(ethoxycarbonyl)-4-(4-fluorophenyl)
thiophen-2-yl)amino)-1,6-dioxohexa-2,4-dien-3-olate (****10****).* Yellow solid (yield 73%),
m.p. 253–254 °С. ^1^H NMR (DMSO-*d*_6_, 400 MHz) δ (ppm): 13.41 (s, 1H), 8.05 (m, 1Н), 7.88 (m, 1Н), 7.72 (m, 1Н), 7.67 (br. s., 1Н), 7.32 (m, 2Н), 7.27 (br. s., 1Н), 7.17 (m, 2Н), 6.93 (s, 1H), 5.98 (s, 1H), 4.16 (q, *J* = 7.0 Hz, 2Н), 1.06 (t, *J* = 7.0 Hz, 3Н). ^13^C NMR (DMSO-*d*_6_, 100 MHz) δ (ppm): 185.6, 164.5, 163.9, 163.9, 162.9, 161.4 (d, *J* = 243.5 Hz), 160.6, 151.0, 139.6, 138.5, 133.9, 133.4 (d, *J* = 2.4 Hz), 131.5, 130.7, 130.4 (d, *J* = 8.1 Hz), 128.7, 127.0, 116.1, 115.5, 114.3 (d, *J* = 21.3 Hz), 114.2, 92.4, 60.0, 13.6. Found, %: C, 52.45; H, 2.83; N, 7.09; S, 5.33. C_26_H_17_Cl_2_FN_3_NaO_5_S. Calculated, %: C, 52.36; H, 2.87; N, 7.05; S, 5.38.
